# Discovery of ongoing selective sweeps within *Anopheles* mosquito populations using deep learning

**DOI:** 10.1101/589069

**Authors:** Alexander T. Xue, Daniel R. Schrider, Andrew D. Kern, Ag1000G Consortium

## Abstract

Identification of partial sweeps, which include both hard and soft sweeps that have not currently reached fixation, provides crucial information about ongoing evolutionary responses. To this end, we introduce *partialS/HIC*, a deep learning method to discover selective sweeps from population genomic data. *partialS/HIC* uses a convolutional neural network for image processing, which is trained with a large suite of summary statistics derived from coalescent simulations incorporating population-specific history, to distinguish between completed versus partial sweeps, hard versus soft sweeps, and regions directly affected by selection versus those merely linked to nearby selective sweeps. We perform several simulation experiments under various demographic scenarios to demonstrate *partialS/HIC*’s performance, which exhibits excellent resolution for detecting partial sweeps. We also apply our classifier to whole genomes from eight mosquito populations sampled across sub-Saharan Africa by the *Anopheles gambiae* 1000 Genomes Consortium, elucidating both continent-wide patterns as well as sweeps unique to specific geographic regions. These populations have experienced intense insecticide exposure over the past two decades, and we observe a strong overrepresentation of sweeps at insecticide resistance loci. Our analysis thus provides a list of candidate adaptive loci that may be relevant to mosquito control efforts. More broadly, our supervised machine learning approach introduces a method to distinguish between completed and partial sweeps, as well as between hard and soft sweeps, under a variety of demographic scenarios. As whole-genome data rapidly accumulate for a greater diversity of organisms, *partialS/HIC* addresses an increasing demand for useful selection scan tools that can track in-progress evolutionary dynamics.

## Introduction

Malaria represents an enormous burden on human health, with an estimated 214 million cases and 438,000 deaths in 2015 [1]. As mosquitos of the *Anopheles gambiae* species complex are the major vector for *Plasmodium* parasites, roughly 70% of global malaria relief budgets have been focused on mosquito control, including insecticide treated bed-nets, indoor residual spraying, and larva control through the direct modification of habitats as well as the application of larvicide. While these vector control efforts have successfully produced major reductions of malaria transmission rates over the past 15 years [1], there has been an alarming increase in mosquitos resistant to insecticides, specifically pyrethroids, observed across nearly all areas of the world covered by anti-malarial efforts [2]. Pyrethroids are the only class of insecticide used in long-lasting insecticidal nets and are applied in many indoor spraying programs, thus the evolutionary innovation of resistance is a well-recognized Achilles heel of anti-malarial efforts.

The increase in insecticide resistant mosquitoes is to be expected from an evolutionary standpoint: anti-malaria control efforts exert a strong selective pressure to which mosquito populations will respond through the differential survivorship and reproduction of those individuals that can best cope with the applied insecticides. Pyrethroid resistance was reported within African malaria vectors first in Sudan during the 1970s, then later in West Africa during the 1990s, most likely stemming from accidental exposure of mosquitos to crop applications of pyrethroids [3,4]. Subsequent analysis showed this earliest resistance to be a result of mutations in the knockdown resistance locus *kdr*, which is known to contribute to pyrethroid resistance in other insect species [5]. Mutations conferring resistance at *kdr* as well as other loci have since spread throughout Africa, and threaten to nullify the gains in malaria control achieved over the past decade [6]. While control efforts are now looking toward non-pyrethroid insecticides [2,7] as well as gene drive technologies [8], it is anticipated that resistance to these control modalities will eventually evolve as well [9,10]. Hence, an important goal in the continued fight against malaria is to identify genomic targets of resistance in *Anopheles*, especially in such a way that might inform vector managers in the field.

Alleles that confer resistance to control efforts should rapidly increase in frequency within *Anopheles* populations in a manner consistent with selective sweeps. When an allele increases in frequency under selection, its linked genetic background comes with it in a process known as genetic hitchhiking. Selective sweeps, through this hitchhiking effect, lead to decreased levels of polymorphism [11–13], skewed allele frequency spectra [14,15], and increases in linkage disequilibrium surrounding the site under selection [16]. Classically, methods for finding sweeps have focused on a particular aspect of genetic variation, for instance observing the site frequency spectrum (SFS) at a locus and comparing it to expectations under neutrality and selective sweeps [17]. More recently, the field has made excellent progress in combining signals across multiple features of genetic variation through supervised machine learning (SML) [18–25], which has substantially improved power, accuracy, and robustness in what have been stubbornly difficult inference problems within population genetics [26]. While much attention has been paid to applying SML for the identification and classification of completed selective sweeps in the genome [22,24], less effort has been made for using SML to identify sweeps that are incomplete within a population, sometimes called partial sweeps (although see Sugden *et al*. [25] for a recent example). In these cases, the beneficial allele is not currently fixed within the population, thereby creating a weaker hitchhiking effect in comparison to a completed sweep, and accordingly a more subtle perturbation of patterns of genetic variation [27]. Although difficult to detect, partial sweeps could be implicated in cases where recently initiated selective forces cause presently ongoing adaptation, directional selection ceases prior to fixation, or an intermediate allele frequency is favored by balancing, polygenic, and/or pleiotropic selection. Therefore, it is important to address the challenge of capturing such genomic signatures that represent a significant facet of evolution, which may also give insight into future dynamics.

## New Approaches

Recent successful efforts to reduce malaria transmission are in danger of collapse due to evolving insecticide resistance in the mosquito vector *Anopheles gambiae*. We aim to understand the genetic basis of current adaptation to vector control efforts by deploying a novel method that can classify multiple categories of selective sweeps, including partial sweep classes, from population genomic data. To this end, we extend a recent SML method to partial sweep inference and apply it to elucidate ongoing selective sweeps from *Anopheles* population genomic samples.

Specifically, we introduce here an extension of *S/HIC* [22] and *diploS/HIC* [24], *partialS/HIC*, a SML classifier that uses a deep convolutional neural network (CNN) to classify a genomic window from a set of selection states that includes both hard and soft partial sweeps along with their associated linked classes (*i.e*. regions adjacent to either a partial hard or soft sweep). Importantly, this implementation achieves increased inferential power by utilizing dozens of additional summary statistics, including summaries for the distribution of *iHS* scores [28] as well as derivatives of a recently developed SNP-specific compound statistic called SAFE (selection of allele favored by evolution) [29], under a deep learning framework that involves training convolutional neural networks with coalescent simulations that accommodate demographic history and are converted into spatially-explicit two-dimensional feature vector images. We validate *partialS/HIC*’s performance through extensive simulation-based experiments that were modeled after data from phase I of the *Anopheles gambiae* 1000 genomes project (Ag1000G) [6], with particular emphasis on discovering sweeps currently in progress such as what might result from ongoing vector control efforts (*e.g*. insecticide spraying). Our findings demonstrate that our method is effective for detecting partial sweeps even in the face of complex population size histories such as those found among the Ag1000G samples. Furthermore, for binary classification of selective sweeps when partial sweeps are included, *partialS/HIC* has greater accuracy than *iHS* or SAFE alone as well as two alternative approaches to sweep inference based on the same suite of summary statistics. Subsequently, we apply our method to the empirical Ag1000G data, revealing many partial sweeps as well as completed sweeps from standing genetic variation. Moreover, we find that our sweep candidates are highly enriched for loci that have been previously identified as contributing to insecticide resistance.

## Results

### Coalescent simulations of feature vector images for *partialS/HIC* training

To train *partialS/HIC*, we deployed the program discoal [30] to perform coalescent simulations of completed and partial as well as hard and soft selective sweeps, along with simulations without sweeps, in a manner analogous to Schrider and Kern [22] (Figure S1). Individual simulations were converted into two-dimensional matrices, or feature vector images, built from 89 rows corresponding to different summary statistics, and 11 columns corresponding to adjacent sub-windows. The 89 statistics include, along with 14 that were previously implemented in *S/HIC* and/or *diploS/HIC*, 3 genomic region variants of the SNP-specific *iHS* statistic [28] and 72 derivatives of the SAFE score [29]. *partialS/HIC* is trained to classify genomic segments into one of nine states: unaffected by selection (*i.e*. neutral); containing a completed hard, completed soft, partial hard, or partial soft sweep, respectively; or linked to a completed hard, completed soft, partial hard, or partial soft sweep, respectively. To this end, we defined the four completed hard/soft and partial hard/soft selective sweep states as containing a sweep within the central, focal sub-window (*i.e*. the fifth out of the 11 columns of sub-windows). In contrast, the four classification states involving linkage to a selective sweep were defined as having a sweep of the given type within one of the remaining ten sub-windows. Therefore, every linked selection state was trained from simulations that vary in genetic distance to the sweep target site, such that the total set of simulations all had a sweep occurring in any of the ten non-focal sub-windows. This allowed us to accommodate a range of linked classes that differ in spatial patterns within a genomic window, as well as assess how linkage distance affects misclassification bias during our simulation experiments. Simulations were run for each of eight population size histories corresponding to the empirical Ag1000G population datasets, which were previously inferred as part of the initial data release [6]. These *Anopheles* population datasets from Miles *et al*. [6] are labeled here as AOM (*A. coluzzii* from Angola), BFM (*A. coluzzii* from Burkina Faso), BFS (*A. gambiae* from Burkina Faso), CMS (*A. gambiae* from Cameroon), GAS (*A. gambiae* from Gabon), GNS (*A. gambiae* from Guinea), GWA (*Anopheles* of uncertain species from Guinea-Bissau), and UGS (*A. gambiae* from Uganda).

Heatmaps constructed from median values across simulations reveal expected spatial patterns, such that values immediately flanking a sweep are substantially different than those further from the focal sub-window, while neutral regions display no discernible pattern among sub-windows (Figure S2). Additionally, spatial patterns of statistics differ qualitatively between selection states. These observations are consistent regardless of mosquito population history, suggesting that there is signal within this collection of summary statistics to isolate the location of a sweep to a specific sub-window as well as distinguish among neutral regions and types of selective sweeps.

### Deep learning excels in detecting selective sweeps, including partial hard sweeps

**Figure 1.**
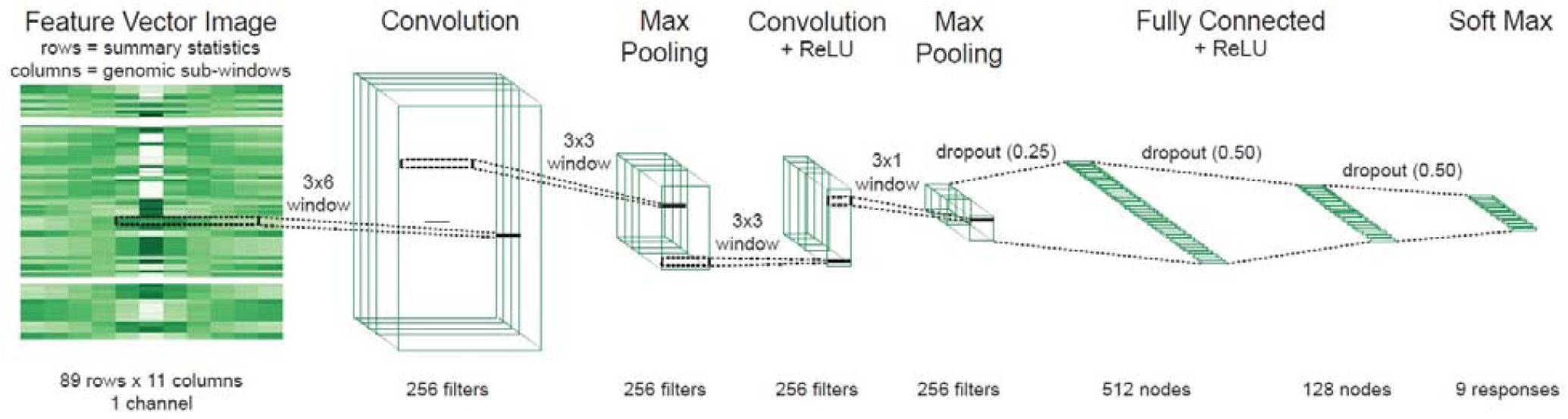
CNN architecture of neural network layers. Our *partialS/HIC* classifier utilizes a convolutional neural network whereby the input feature vector image, composed of 89 summary statistics organized into rows and across 11 contiguous genomic sub-windows organized into columns, is passed to a 2D convolutional layer with 256 filters using a 3×6 receptive field. Next is a 2D max pooling layer given a 3×3 receptive field. Then there is a second 2D convolutional layer of 256 filters based also on a 3×3 receptive field and ReLU activation. Afterward is a second 2D max pooling layer with a 3×1 receptive field. The tensor is then flattened after a dropout layer (p=0.25) and passed to two fully-connected layers with ReLU activation, resulting in 512 and 128 nodes with subsequent dropout (p=0.50) respectively. Finally, a softmax activation layer classifies the 9 responses.

We utilized *partialS/HIC* to train and test a CNN for nine-state classification independently on each of the eight demographic histories associated with the Ag1000G population samples (Figures S1, 1). To this end, we produced eight batches of simulations that were split into separate training, validation, and testing sets. During the training process, CNN hyperparameters were tuned on the training set while the validation set allowed mitigation of over-fitting (Figure S3). Each CNN was subsequently assessed for accuracy with the corresponding held-out testing set, which was generated under the same specifications as the training/validation data except linked selection classes were kept discrete versus being grouped together into the four linked selection states. Among the eight test sets, there was moderate overall accuracy for this simulation experiment (median accuracy = 66.4%; Table S1). However, confusion matrix heatmaps provide a more informative view of our classifier’s performance, which exhibited reliability in identifying neutral regions, completed hard and soft sweeps, partial hard sweeps, and individual regions linked to completed hard/completed soft/partial hard sweeps (Figures 2, S4). In general (*i.e*. all demographic scenarios save for AOM), completed hard sweep is the class that experienced the highest degree of correct assignment (median accuracy = 96.0%). We also had high accuracy for identification of linked completed hard regions, demonstrating a strong ability to localize completed hard sweeps. The behavior of our *partialS/HIC* classifier is likewise favorable for completed soft sweeps (median accuracy = 84.2%), with completed soft sweeps rarely detected incorrectly as hard sweeps and instead typically mistaken for either the neutral or partial soft sweep state. Moreover, sub-windows linked to completed soft sweeps beyond one sub-window away had low levels of misclassification to any of the non-linked sweep states, again allowing for dependable localization of the sweep.

**Figure 2.**
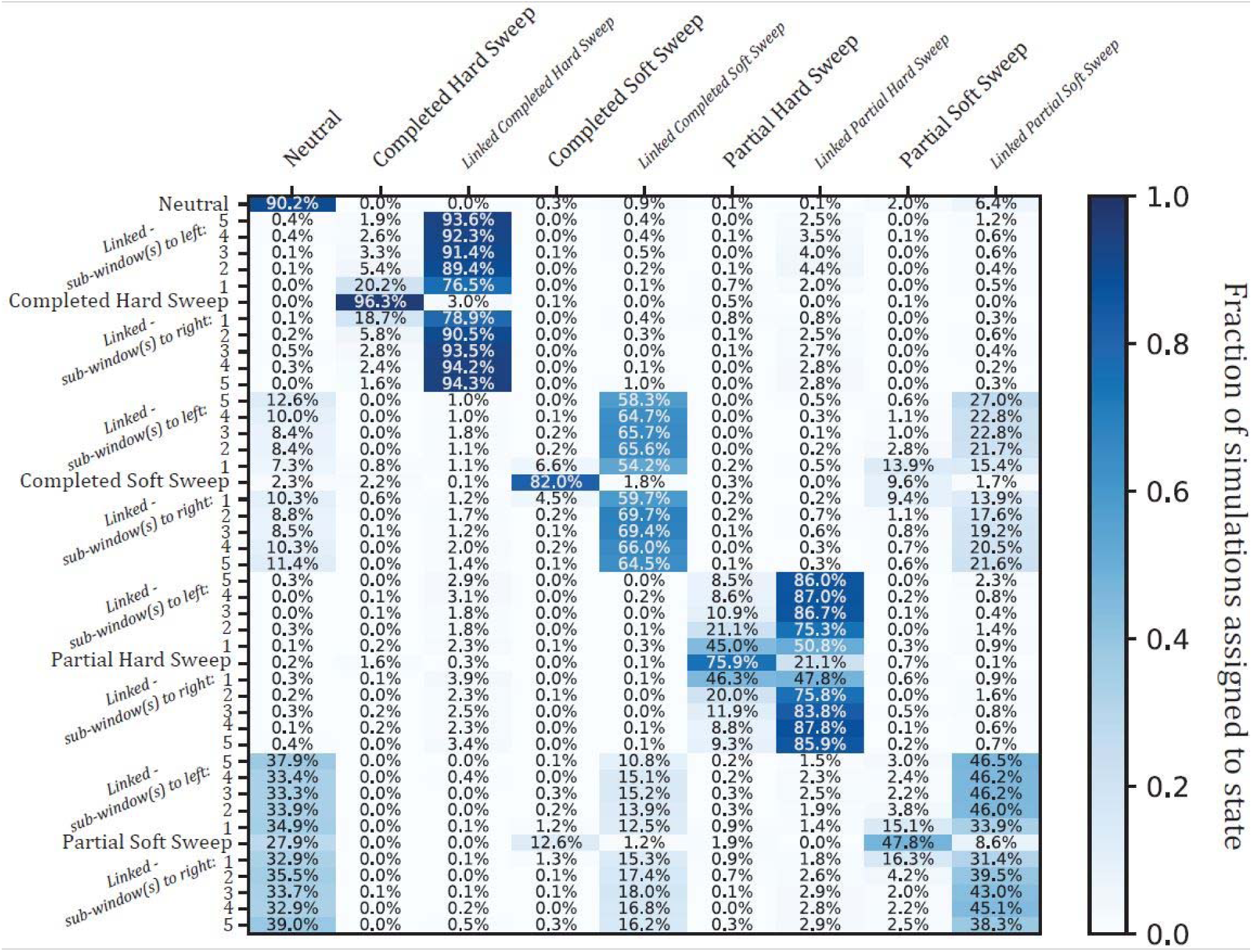
Confusion matrix heatmap of *partialS/HIC* simulation experiment. Given the BFS population history. Each row designates a true simulation class, which for linked sweeps is differentiated by distance in genomic sub-windows to the target selective sweep. In total, there are 45 simulated scenarios shown, including 11 for each sweep type (i.e. one case whereby the sweep is within the central sub-window, and 10 whereby a linked sweep is located within one of the flanking sub-windows) and neutrality. Each column indicates one of nine inferred states, with the linked simulation classes collapsed into a single category per selective sweep type for training and classification, allowing *partialS/HIC* to learn to distinguish between sweeps located in the focal sub-window and those somewhere nearby. Darker blue cells represent a higher proportion of the 1,000 calls for each true class. Importantly, there is generally high accuracy in discriminating between sweeps within the focal sub-window versus linked sub-windows, especially when linkage is beyond one sub-window away and particularly for completed sweeps. However, discovering partial soft sweeps is noticeably a challenging task. Moreover, there is greater sensitivity separating full completion from partial completion for hard sweeps in contrast to soft sweeps.

Importantly, the purpose of *partialS/HIC* is to extend our state space to identify ongoing selective sweeps while distinguishing these from completed sweeps. We find that our ability to identify partial hard sweeps was generally strong across population histories (median accuracy = 74.6%) and is often comparable to that of completed soft sweeps. However, localization of partial hard sweeps along the chromosome was more difficult than for completed sweeps, as can be seen from the moderate levels of confusion between partial hard sweep and linked partial hard sweep sub-windows. Undoubtedly, this is due to the limited amount of time recombination has had to whittle down the haplotype carrying the beneficial mutation.

Identifying partial soft sweeps was a much more challenging task (median accuracy = 45.3%), with a high false negative rate (median rate of misclassification as neutral = 27.6%) as well as a substantial probability of misclassification as a completed soft sweep (median rate = 14.4%). It is encouraging though that partial soft sweeps were almost never misclassified as a completed nor partial hard sweep. Additionally, while our accuracy in classifying partial soft sweeps was poor, false positives were not a major concern (median rate of misclassifying neutral regions as partial soft sweep = 3.2%). Although even fairly low false positive rates can be problematic when the true number of sweeps is low, *partialS/HIC* achieved acceptably low false discovery rates for partial soft sweeps in our application to the *Anopheles gambiae* 1000 Genomes dataset, as we show below. Furthermore, linked partial soft sweeps beyond one sub-window away from the focal sub-window were rarely mistaken for a sweep state, and likewise partial soft sweeps were seldom confused for being linked (median rate of misclassification as linked partial soft sweep = 4.3%), thus demonstrating that accurate localization of partial soft sweeps may be possible.

In summary, *partialS/HIC* has excellent ability to distinguish partial from completed sweeps for *de novo* mutations, and lesser yet still substantial power for sweeps from standing variation. Moreover, we demonstrated very strong performance in differentiating between hard and soft sweeps, regardless of whether a sweep was completed or incomplete. Importantly, this is all while maintaining an acceptable false positive rate across each of the population histories tested (median accuracy for neutral regions = 85.1%; median rate of misclassifying neutral regions as any one of the four non-linked sweep states = 4.6%).

### Robustness to demographic model misspecification

To assess robustness to demographic misspecification, we applied a CNN trained on simulations from one population sample to data generated from an alternate demographic history. Specifically, we used training data from the GAS population size history, which was fairly stable over time, and leveraged it against the CMS test dataset, which experienced a dramatic population expansion (overall accuracy = 55.0%; rate of misclassifying neutral regions as any one of the four non-linked sweep states = 1.7%). Despite this misspecification, the confusion matrix (Figure S5) strongly resembles the corresponding matrix that is correctly specified for demography (*i.e*. for CMS in Figure S4). In particular, accuracies for finding neutral regions, completed hard sweeps, and partial hard sweeps are roughly equivalent between the correctly specified model and misspecified model (Figure S5). For soft sweeps, while confusion between completed and partial sweeps is increased for the misspecified model, the overall ability to distinguish sweeps from neutrality is largely preserved. Moreover, the rates at which examples from the linked classes were mistaken for sweeps are seemingly unaffected.

To further examine the impact of increasingly misspecified demography, we produced five additional full testing datasets assuming a simple two-epoch instantaneous contraction model. The five test sets differed only by bottleneck severity, which increased in even intervals from 20× to 100×. We then applied the CNN that was trained given the BFS population history and measured overall accuracies. These accuracy measurements are generally similar to that of our baseline, *i.e*. the corresponding original experiment whereby the testing data were simulated under the inferred demography for BFS, thus the demography was correctly specified with respect to the training set underlying this CNN (Figure S6). However, performance gradually worsens with decreasing bottleneck severity across the test sets, but this may be largely an effect from the number of polymorphisms. To address this, we generated another six test sets that identically replicated the specifications for the BFS training simulations except with *θ* fixed in value yet varying between the six datasets. Notably, the range we utilized here for *θ* exceeded the *θ* prior distribution for the original BFS training and testing simulations, hence further evaluating model misspecification. We find no loss in overall accuracy except in the case with the least genetic diversity simulated, where there is a moderate decrease (Figure S7). This is unsurprising given that low levels of genetic diversity lead to noisy estimates of the statistics employed by *partialS/HIC*. It is encouraging however that sensitivity is maintained throughout most of our *θ* prior distribution range and accuracy does not fall below that of our baseline experiment until *θ* is at a low value toward the bounds (*θ* < 5,000 for the full genomic window size of 55,000 base-pairs). To put this into perspective, genetic diversity measures for the corresponding empirical BFS dataset are consistent with simulations generated with *θ* ≥ 5,000, and likewise are beyond nearly the entirety of the *θ* = 1,000 simulated values; among the sub-windows that passed our filtering thresholds (Figure S1), there is a median value of *θ*_*W*_ = 2.633e-2 (central 95% density: 6.484e-3 – 3.547e-2) and *θ*_*H*_ = 6.318e-3 (central 95% density: 2.381e-3 – 1.231e-2), compared to a median value of *θ*_*W*_ = 1.218e-2 (central 95% density: 1.116e-3 – 1.849e-2) and *θ*_*H*_ = 3.863e-3 (central 95% density: 8.592e-4 – 9.696e-3) across all sub-windows generated under the *θ* = 5,000 condition, whereas the simulations given *θ* = 1,000 have a median value of *θ*_*W*_ = 2.353e-3 (central 95% density: 9.802e-5 – 3.861e-3) and *θ*_*H*_ = 7.043e-4 (central 95% density: 5.950e-8 – 2.571e-3).

Together, these results indicate that sweep discovery and localization is not strongly impacted by several forms of demographic model misspecification during training.

### Partial sweeps are unpredictably misclassified as either completed sweep or neutral when not explicitly considered

Since the previous versions of *partialS/HIC* (*S/HIC* and *diploS/HIC*) did not allow for partial sweep selection states, we were interested in how such five-state classifiers would behave when confronted with partial sweeps. To explore this, we conducted a simulation experiment that first removed partial hard and soft sweeps as well as their associated linked classes from the CNN training process, thus training on only five states rather than all nine. Next, in an effort to examine the classification behavior for these five-state CNNs, we applied the full test set that included the partial sweep classes. Unsurprisingly, the trend was for partial sweeps to be most often confused for linked selection (Figures 3, S8). Perhaps more concerning is the false negative rate (*i.e*. rate at which partial sweeps were misclassified as neutral), which was substantial in partial hard sweeps for several populations (median = 8.8%; max = 32.4%; >1% in all populations) and extreme in partial soft sweeps (>50% in three populations, >40% in three more populations, and >24% in all populations). Partial hard sweeps that were discovered were also often misclassified as a completed soft sweep (median rate of misclassification as completed soft sweep = 5.5%). However, when training included partial sweeps, there is universal and dramatic improvement in both finding sweeps and correctly identifying the model of selection (Figures 2, S4). Meanwhile, overall accuracy remains similar among the five-state and nine-state classifiers with respect to simulations of neutral, completed sweep, and linked classes exclusively (Figures 3, S8; Table S2). As a result, accuracy should only stand to benefit from incorporating partial sweeps into training since ignoring such information leads to unacceptably high false negative rates of partial sweeps being called neutral or linked.

**Figure 3.**
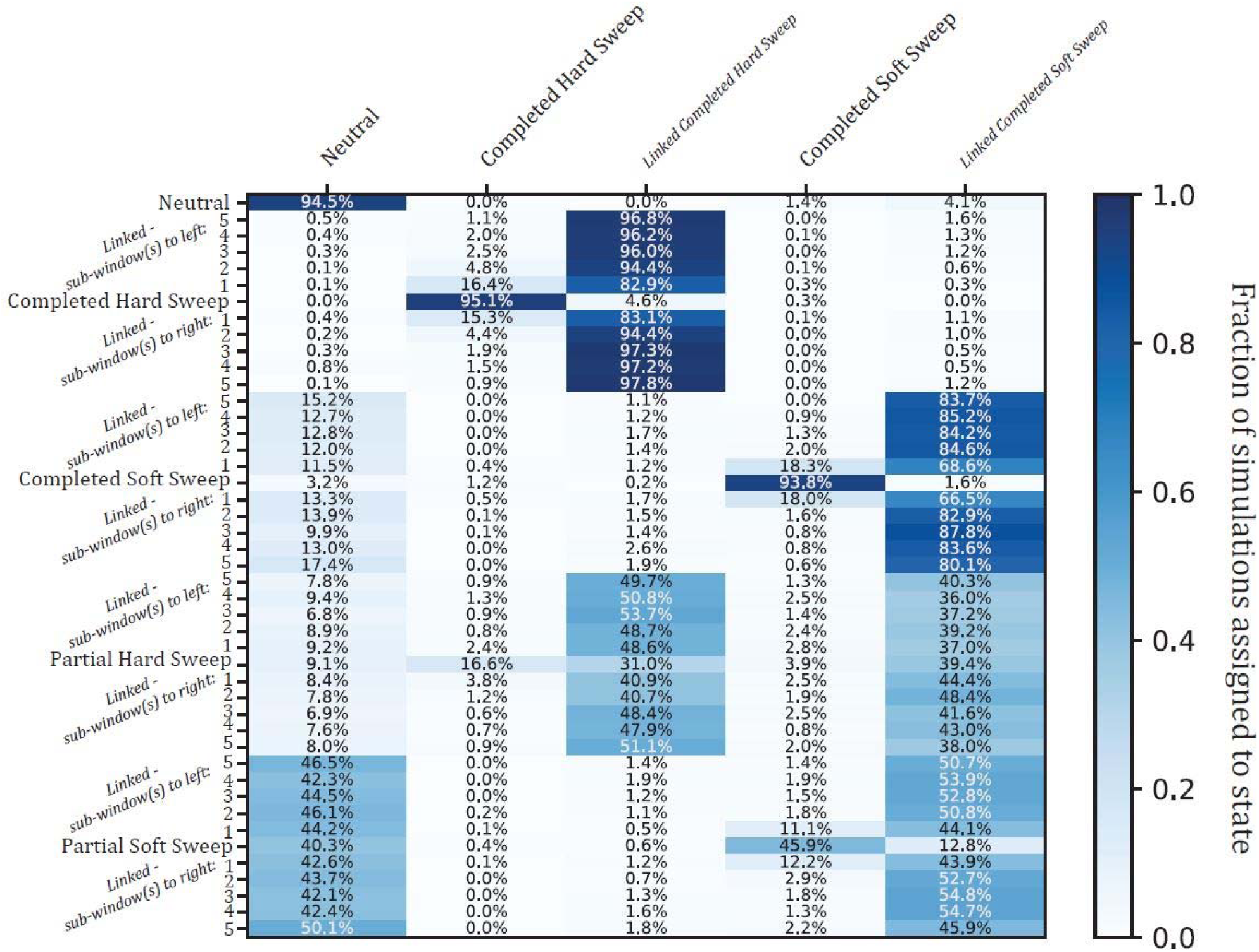
Confusion matrix heatmap of simulation experiment with partial sweeps ignored during training. Given the BFS population history. Structured in the same manner as Figure 2, each row designates a true simulation class and darker blue cells represent a higher proportion of the 1,000 calls for each true class. In contrast to Figure 2 though, each column indicates one of five inferred states instead of nine, as the two partial sweep and two linked partial sweep classification states were omitted from training to determine misclassification bias when partial sweeps are ignored. There is a substantial decrease in discovery of partial sweeps, especially partial hard sweeps (20.5% total sweep discovery compared to 78.2% in Figure 2; 46.3% total sweep discovery of partial soft sweeps compared to 62.3% in Figure 2). Specifically for partial hard sweeps, a large proportion of the detected sweeps are misclassified as soft sweeps instead of hard sweeps (19.0%; i.e. 3.9% of calls = soft sweeps: 16.6% of calls = hard sweeps).

### *partialS/HIC* binary classification outperforms several competing approaches

To assess whether the collection of summary statistics under our deep learning method extends inferential resolution beyond the signal conferred by *iHS* or SAFE alone, and to determine if our CNN approach leverages these statistics in a more informative manner than other aggregation schemes, we compared receiver operating characteristic (ROC) curves, which plot true positive against false positive rates given varying thresholds, for the binary classification task of broadly detecting selective sweeps (*i.e*. any of the four selection states involving a sweep within the focal sub-window) vs. neutral regions or linked sweeps. We selected for our comparison *iHS*-derived statistics (mean, maximum, and proportion of outlier *iHS* values within the central sub-window of the testing simulations) because *iHS* was explicitly designed for detecting partial sweeps [28], and SAFE-based statistics (variance and maximum of SAFE scores within the central sub-window of the testing simulations) since SAFE is a compound of many variables, of which the vast majority of our training data are summaries [29]. Additionally, we conducted a principal component analysis (PCA) from the training data, restricted to only the focal sub-window as well as for the entire dataset throughout the full window respectively in two separate PCAs, and projected the testing data onto the first two principal components (PC1, PC2). We then produced a ROC curve from each set of principal component values transformed from the testing data, respectively to both PC1 and PC2 as well as the PCA on solely the central sub-window versus the full-scale window, for a total of four individual ROC curves. These ROC curves allow contrast to a simple dimensionality reduction approach. Lastly, we computed Composite of Multiple Signals scores from the testing data for both the focal sub-window and full window respectively, using the training data to derive probability distributions. Deploying the Composite of Multiple Signals metric provides an alternative method that is intended to exploit multiple summary statistics for inference of positive selection [31]. For a direct comparison, we altered *partialS/HIC* slightly to create a CNN that had two final output responses (*i.e*. sweep in central sub-window vs. no sweep in central sub-window) instead of nine, with the training dataset as well as architecture of network layers (Figure 1) remaining the same. The CNN optimized for binary classification was then applied to the testing simulations, with the output probability of a selective sweep utilized as input to generate the ROC curve.

The *partialS/HIC* binary classifier consistently outperforms other methods in identifying selective sweeps to the focal sub-window when partial sweeps are included (median AUC = 0.943; Figures 4, S9). Interestingly, we demonstrate that the Composite of Multiple Signals also has a strong ability to extract information from our selection of summary statistics, though it always trails behind *partialS/HIC*; in every case, the best performance after *partialS/HIC* is for the Composite of Multiple Signals based on all sub-window statistics (median AUC = 0.876), while the next best ROC curve is from the Composite of Multiple Signals calculated from only the central sub-window (median AUC = 0.859). In contrast, with the unexpected exception of PC2 for central sub-window statistics (median AUC = 0.749), PCA appears to be an unfavorable approach for combining summary statistics to infer selective sweeps (*e.g*. median AUC for PC1 of all sub-window statistics = 0.500). Furthermore, there is evidence that SAFE scores capture important signal about selective sweeps in a manner that is robust to the final frequency of the selected mutation, particularly the variance of SAFE scores within the focal sub-window (median AUC = 0.787). Conversely, there seems to be decreased signal in *iHS*, as all its focal sub-window variants individually performed quite poorly (*e.g*. median AUC for proportion of *iHS* outliers = 0.582).

**Figure 4.**
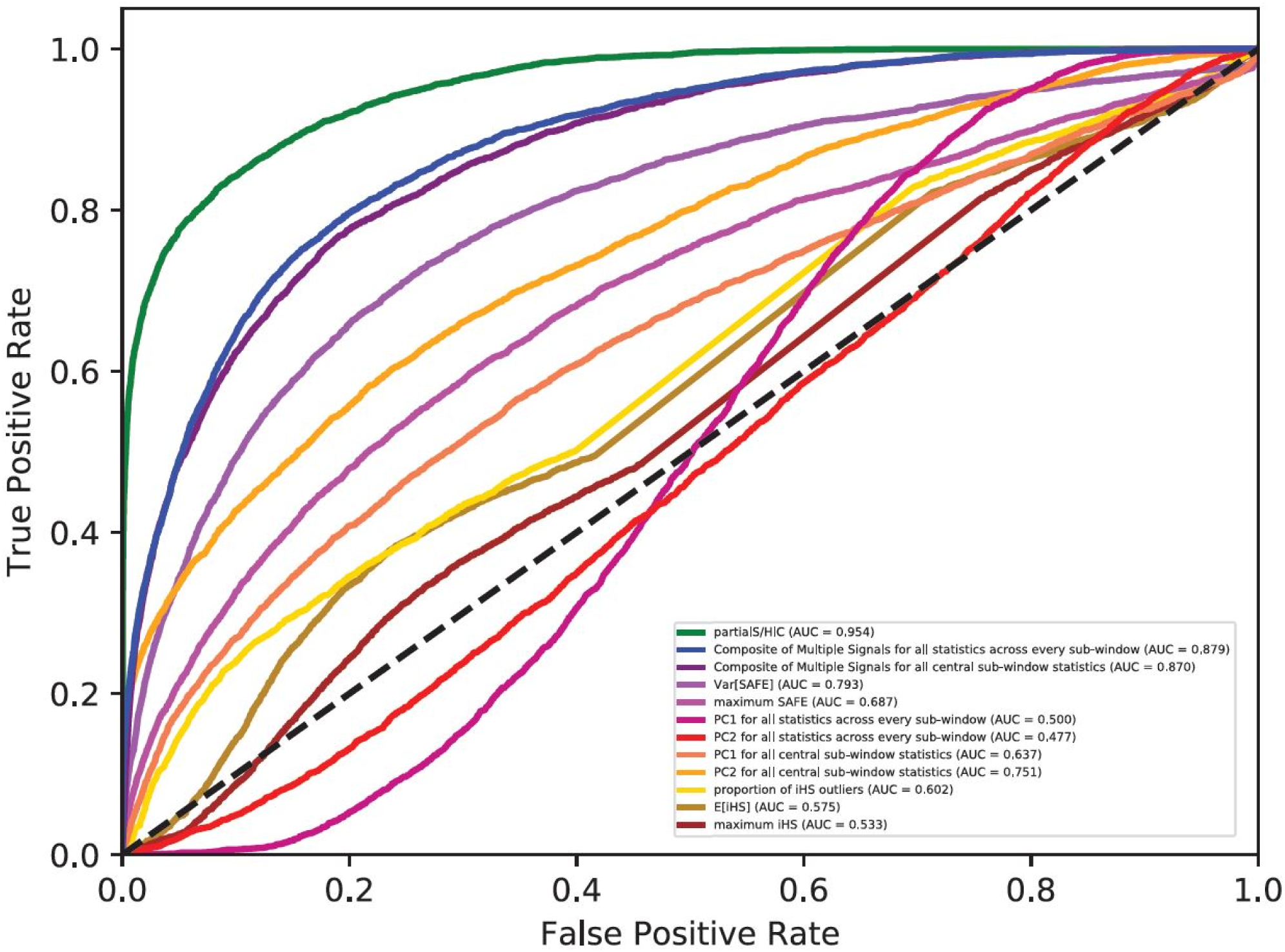
ROC curves for binary classification of selective sweeps, including partial sweeps, versus regions neutrally evolving or under linked selection. Given the BFS population history. The *partialS/HIC* deep learning classifier outcompetes two other approaches for managing the same suite of summary statistics: Composite of Multiple Signals and principal component analysis. Furthermore, *partialS/HIC* excels in performance over several sub-window derivatives of two summary statistics, SAFE and iHS, which were all included within our set of summary statistics used for training. Notably, Composite of Multiple Signals, SAFE, and iHS were all designed to uncover selective sweeps. Additionally, the SAFE score is itself a compound statistic that captures signal from several constituent statistics, and the majority of our training data is derived from either the SAFE score or one of its components.

### Soft and partial sweeps are commonplace among *A. gambiae* populations

Turning our attention to the Ag1000G phase I data, we applied our nine-state CNNs to the corresponding *A. gambiae* population datasets, classifying 5 KB segments using a 55 KB full sliding window throughout the whole genome (Figure S1). Each of the eight mosquito populations contains a large number of sub-windows identified as completed soft sweeps (median fraction of total calls genome-wide = 5.01%) as well as partial sweeps (median fraction of total calls genome-wide for partial hard sweep = 2.84%; median fraction of total calls genome-wide for partial soft sweep = 7.24%), coupled with only a handful of completed hard sweep predictions (median fraction of total calls genome-wide = 0.03%) (Figure 5; Table S3). Partial soft sweeps were typically discovered most often (median proportion of sweep calls = 52.59%), with completed soft sweeps often following (median proportion of sweep calls = 28.80%) and partial hard sweeps usually being the third most numerous class of detected sweep (median proportion of sweep calls = 19.75%). Notably, our estimated false discovery rates (FDRs) are higher for soft sweeps (median FDR for completed soft sweeps = 11.09%; median FDR for partial soft sweeps = 12.20%) compared to hard sweeps (median FDR for completed hard sweeps = 0.00%; median FDR for partial hard sweeps = 0.39%); this implies that individual soft sweep candidates should be viewed with more caution, though we should be able to estimate the genome-wide proportion of these classes well. Specifically, classifications for partial soft sweeps outnumber those for completed soft sweeps in every population besides GNS, as well as partial hard sweep calls in all populations but GAS. Importantly, these findings remain the same, with the exception that calls for partial hard sweeps now slightly exceed partial soft sweep classifications in GNS, after false discovery correction (Table S3). Additionally, had partial sweeps not been accounted for in the training process, our results suggest that we would have both underestimated the total number of sweeps and incorrectly labeled many of our partial sweeps (Figures 3, S8). This would have led to the conclusion that adaptation from standing variation rather than *de novo* mutations dominate selective sweep dynamics in these *A. gambiae* populations. While it is clear that soft sweeps are indeed more common in these data, our results suggest that hard sweeps often occur as well, though with few reaching fixation. Furthermore, the *partialS/HIC* classifications indicate that most selective sweeps in these population samples are incomplete, suggesting that we are capturing a view of selection in progress.

**Figure 5.**
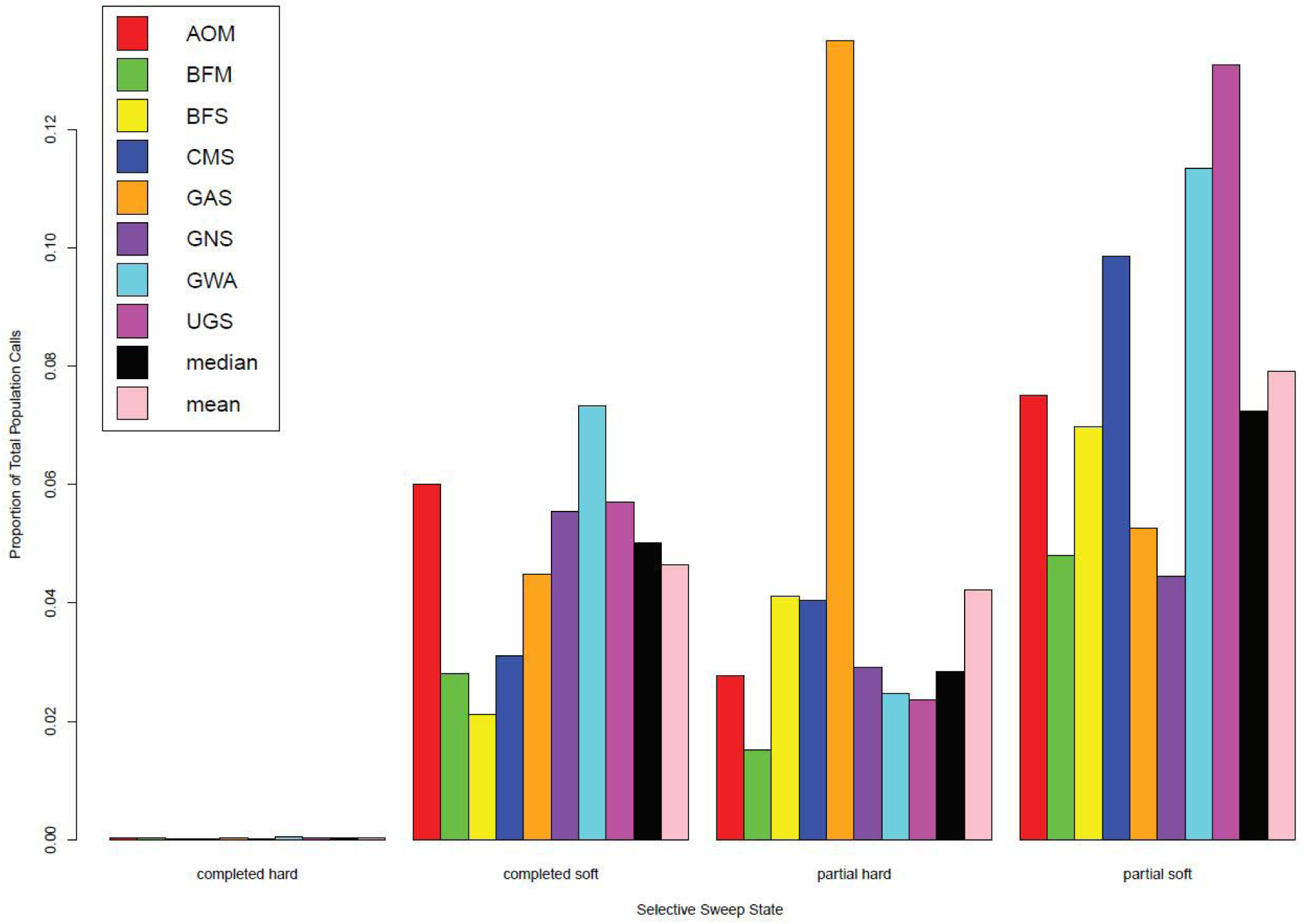
Genome-wide *partialS/HIC* sweep calls across empirical mosquito population datasets. For all eight datasets, there is a minimal number of calls for completed hard sweeps, represented here on the Y-axis as the relative proportion to the total set of empirical calls across every genomic window per population dataset. However, there is indeed a substantial proportion of sweep calls for hard sweeps, yet are incomplete or in-progress. In most cases, the number of both completed and partial soft sweeps further exceeds that of partial hard sweeps. Furthermore, the majority of sweeps have not reach fixation across these Anopheles populations.

### Selective sweeps are significantly enriched in functional regions of the *A. gambiae* genome

To elucidate broad characteristics underlying the genomic targets of selection, we used permutation tests of sweep call locations to discover enrichment patterns in the following DNA regions of interest: gene, mRNA, exon, CDS, five-prime UTR, and three-prime UTR. Permutation tests were based on the total number of calls for the four selection states with sweeps occurring within the focal sub-window, as well as the individual number of calls for each of these states respectively. Across all eight population datasets and for all six DNA regions under investigation, there is a statistically significant enrichment of total sweep calls along with completed soft sweeps calls in particular, whereas completed hard sweep calls are not significantly enriched in any single case (Figures 6, S10; Table S4). Conversely, partial sweep enrichment varies among populations as well as individual DNA regions. Specifically, partial hard sweeps are significantly enriched in five of the six DNA regions for BFS and CMS, and four of the DNA regions for UGS; while partial soft sweeps are significantly enriched in all six DNA regions for UGS, five of the DNA regions for BFM, and four of the DNA regions for BFS.

**Figure 6.**
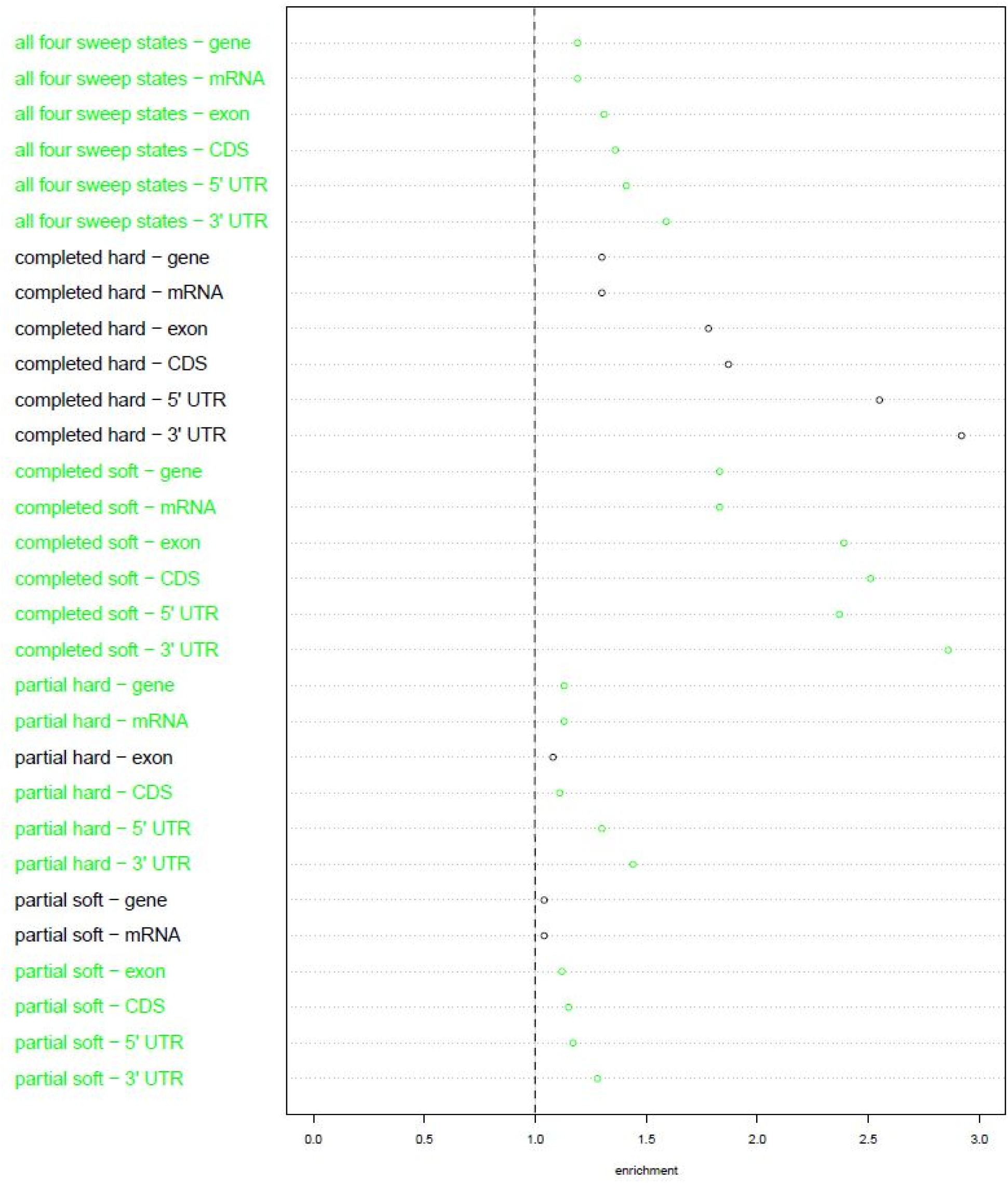
Enrichments of *partialS/HIC* empirical selective sweep calls within DNA regions. For the BFS population empirical dataset. Statistically significant (p < 0.05) enrichments in green. No enrichment (i.e. single-fold enrichment based on the permuted calls) is represented by the dotted line. Completed soft sweeps are enriched within all six DNA regions that experienced permutation tests. Similarly, all of the DNA regions have an enrichment of partial sweeps, hard and/or soft. In contrast, there is no significant enrichment of completed hard sweeps for any of the DNA regions. Overall though, selective sweeps in general are enriched for each of the DNA regions.

### Insecticide resistance loci, especially related to metabolism, are significantly enriched for selective sweeps

We performed a similar permutation analysis for four sets of genes known to confer insecticide resistance (IR), finding at least one set of IR genes to be statistically significant for every population dataset in enrichment of total sweep calls (*i.e*. aggregate of all four sweep classes) and completed soft sweeps, respectively (Figures 7, S11; Table S5). In particular, metabolism-related IR genes are significantly enriched for each of these cases. Furthermore, IR genes corresponding to well-characterized resistance loci (*i.e*. target sites) are significantly enriched for AOM and BFM total sweep calls as well as completed soft sweeps in AOM, BFM, and GNS; IR genes associated with behavior are significantly enriched for BFM and GNS total sweep calls as well as completed soft sweeps in AOM, CMS, and GAS; and IR genes affiliated with cuticular activity are significantly enriched for completed soft sweeps in GWA and UGS. In contrast, completed hard sweeps are only significantly enriched in BFS, GNS, GWA, and UGS for IR genes connected to behavior (as well as metabolism for BFS). For partial sweeps, significant enrichment only occurs within BFM (partial soft sweeps in metabolism as well as behavior IR genes), CMS (partial hard sweeps in metabolism as well as target site IR genes), and UGS (partial hard sweeps in metabolism IR genes).

**Figure 7.**
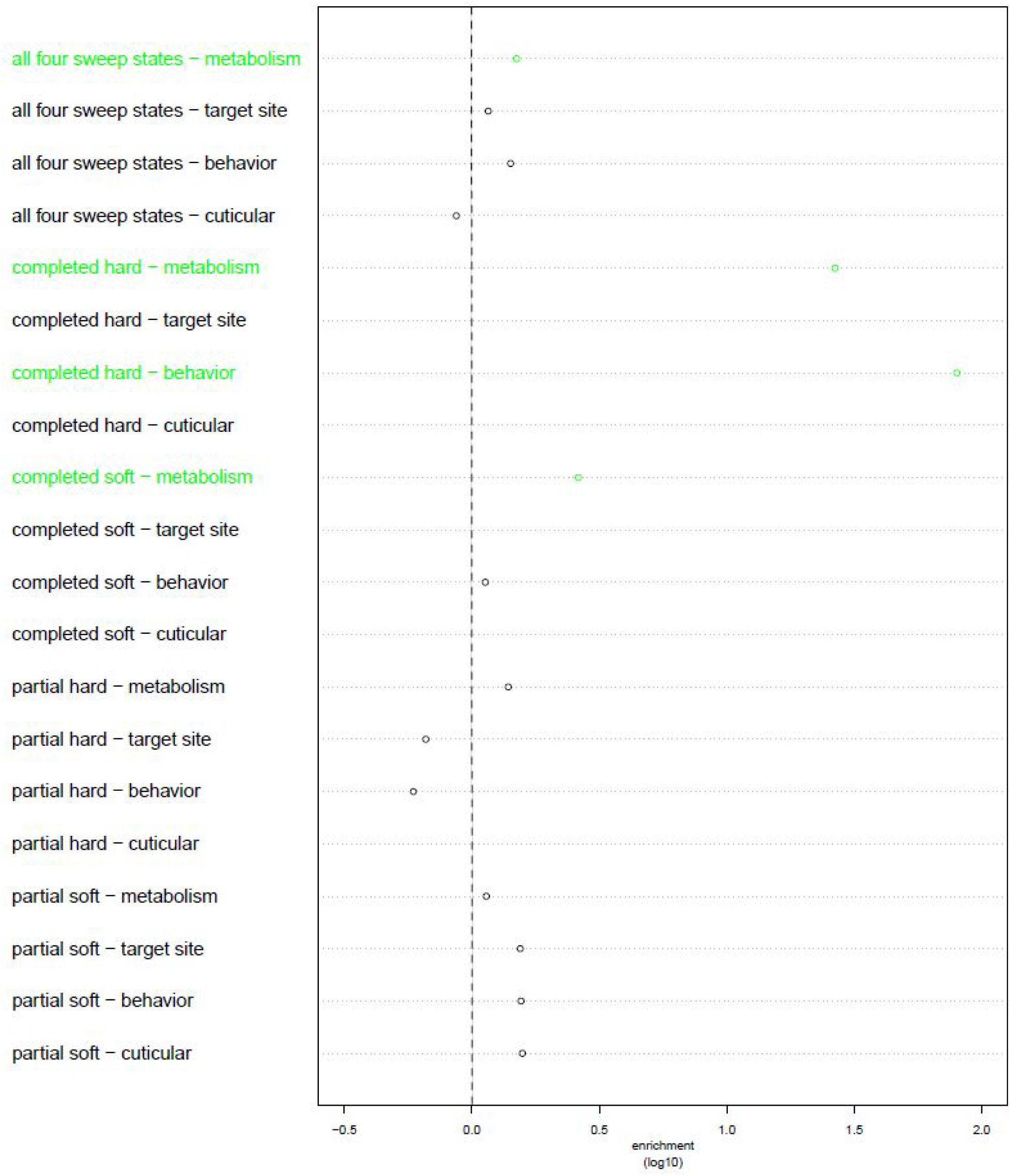
Enrichments of *partialS/HIC* empirical selective sweep calls within groupings of insecticide resistance genes. For the BFS population empirical dataset. Statistically significant (p < 0.05) enrichments in green. No enrichment (i.e. single-fold enrichment based on the permuted calls) is represented by the dotted line. Rows without a plotted point are due to the insecticide resistance gene group not having any calls of that sweep type (e.g. there are no completed hard sweeps called within target site genes). In this mosquito population, there is evidence for several mechanisms through which insecticide resistance evolved, both in sweep types and categories of genes under selection.

### Completed soft sweeps are significantly enriched within the same GO term annotations across populations

To uncover further functional traits targeted by selection, we used our permuted datasets to find individual gene ontology (GO) terms that are enriched in our sweep candidates. Partial soft sweeps are not significantly enriched for any GO terms among populations, while completed hard sweeps and partial hard sweeps are each enriched only in UGS for a single GO term belonging to the cellular components GO domain; specifically, “membrane” is enriched of completed hard sweeps and “nuclear cohesion complex” is enriched of partial hard sweeps. Otherwise, GO term enrichment pertains to just completed soft sweeps as well as the total of all four sweep states in combination (Table S6). For the completed soft sweep significant enrichments, we found six cases of the same GO term in all eight populations. Three of these are related to cellular components (“nucleus”, “membrane” and “integral component of membrane”), and the other three are connected to molecular function, specifically binding (“nucleic acid binding”, “protein binding” and “ATP binding”). All six of these terms, especially those conferring binding function, are also enriched for total sweep calls across multiple datasets: “nucleic acid binding” and “ATP binding” in seven populations; “protein binding” and “nucleus” in six populations; “integral component of membrane” in five populations; and “membrane” in three populations, one of which is significantly enriched for this GO term of completed hard sweeps as well, the single example of such among all populations. Other cases involving the same GO term significantly enriched for completed soft sweeps in over half of the populations include: “binding”, “cytoplasm”, and “zinc ion binding” in seven populations; “RNA binding” in six populations; and “mRNA splicing, via spliceosome” and “ATPase activity” in five populations.

## Discussion

### *partialS/HIC* elucidates both species-wide and population-specific sweep dynamics within *A. gambiae*

The Ag1000G data provided the opportunity to investigate selection at both the continental scale, where wide-reaching impact across the whole species complex could be uncovered, and the regional level, revealing population-specific sweep dynamics. For the former, we observed that *A. gambiae* populations consistently experienced very few completed hard sweeps, with nearly all sweeps being partial and/or soft. In fact, the impact of completed hard sweeps on the adaptive process within mosquitos appears to be even more limited than what was observed previously in humans [32]. This is likely a result of the much larger population sizes and concordant levels of genetic variation that are maintained within *Anopheles* populations. Importantly, we find a large number of ongoing selective sweeps within these populations, particularly in comparison to the number of completed sweeps. There are multiple reasons why this might be the case. A trivial explanation may simply be that we only have power to detect sweeps that have completed in the past few hundred generations, though this seems unlikely. More plausibly, a large number of ongoing sweeps might be expected given the recent change in environment induced by vector control efforts. Another possible explanation is that the frequency dynamics of beneficial alleles within a population is often more complex than assumed and may indeed contain an overdominant component [33]. This would mean that some portion of the partial sweeps that we are observing in *Anopheles* is actually balanced, or transiently balanced, polymorphisms. A fourth class of explanation is that beneficial mutations may not be able to fix in populations due to competition with beneficial mutations on other genes that have originated in different parts of the species range [34]. Indeed, each of these factors may play some role in our reported abundance of partial sweeps.

Although such genome-wide sweep patterns occur species-wide, enrichment behavior seems much more population-specific. For instance, while every population possesses significant enrichment of completed soft sweeps coupled with no completed hard sweep significant enrichments for the six functional DNA regions studied here, partial sweep enrichments vary widely among datasets. Sweep behavior is even more idiosyncratic for insecticide resistance genes, as the only constant between populations is that metabolism is a recurring target of selection, especially for completed soft sweeps.

These findings from the Ag1000G data provide important genomic resources that could inform continental-wide malaria control strategies for the entire *A. gambiae* species complex, as well as have relevance to management efforts specialized to certain populations and localities. Such insight into mosquito vector evolution may also help curb future insecticide resistance adaptation, and in turn prevent impending crises of vector control failure. However, it is important to consider that our partial sweep calls could be capturing more complex selective dynamics at play, for example polygenic and quantitative trait adaptation [35,36], balancing selection [37], and introgression of beneficial alleles. These could lead to different modes of adaptation for the same genomic region across populations, for instance a favorable SNP undergoing a soft sweep at its origin and then carried to neighboring populations (as was suggested in Miles *et al*. [6]) may appear to be experiencing a partial hard sweep in those recipient populations. Such complicated interactions merit further investigation on the Ag1000G data, which would not only continue advancing methodological development for population genetics, but also address interesting questions for a widespread and ecologically important organism that has crucial ramifications on wildlife management and public health.

### *partialS/HIC* offers unprecedented detection of partial sweeps

Supervised machine learning approaches are rapidly gaining traction among population geneticists, with deep learning in particular beginning to experience increased attention and methodological development due to its exciting potential to unlock classic population genetics problems. Examples of successful SML implementation in population genomics include demographic model choice [38], demographic parameter inference [39], comparative analysis of independent single-population size changes [40], identification of introgressed regions [41], recombination rate estimation [42–44], and genomic scans of selective sweeps [22]; deep learning specifically has been employed for joint inference of demography and selection [23], discovery of recombination hotspots [45], estimation of demographic and recombination parameters [46,47], discovery of functional variants [48], prediction of geographic origin [49], and differentiating between hard and soft sweeps from neutral regions [24]. These applications especially benefit from the ability to handle high dimensional input data and bypassing the need of a likelihood function, which is due to SML uncovering data patterns from leveraging *a priori* information through a training algorithm [23,26]. CNNs expand this utility to image processing, which has been demonstrated with *diploS/HIC* to be a powerful tool for exploiting the genomic spatial distribution of multiple population-level summary statistics to detect selective sweeps [24].

Here, we demonstrated with *partialS/HIC* that deep learning can be extended to partial sweeps, especially partial hard sweeps, yielding greater accuracy and robustness than has been previously attained. We also showcased consistent performance in the face of several underlying demographic backgrounds. Specifically, *partialS/HIC* is capable of discovering selective sweeps when partial sweeps are considered, simultaneous disambiguation between partial and completed sweeps as well as between hard and soft sweeps, and spatial localization of selection targets in the genome. Moreover, we have shown that partial sweeps remain mostly undetected if ignored from the training process, even though such selection may be commonplace throughout a genome as with the Ag1000G data. As a result, many previous studies scanning for either complete or ongoing selective sweeps solely (*i.e*. not jointly inferring both types of selection) may have overlooked an important subset of evolutionary events [34]. Researchers may then be interested in reexamining datasets with *partialS/HIC* to elucidate the relative contributions of fixed versus incomplete sweeps to adaptive evolution.

Importantly, the efficacy of *partialS/HIC* relies on several factors that are unexplored here, including simulation prior specifications, CNN architecture with respect to construction and parameterization of neural network layers, and data structure. Hence, it is prudent for future implementations to validate performance by testing a range of configurations, given a project’s individual intricacies, to assess robustness and inherent assumptions. In particular, future exploration of alternate image constructions could be potentially of great methodological benefit. Such images could be derived from different ordering schemes and/or suites of summary statistics, as well as without summary statistics entirely, instead directly exploiting sequence alignments [45–47,49] or even raw reads. More broadly, CNNs can be further extended to address other long-standing efforts in evolutionary biology, such as parameter inference under complex isolation-migration models or phylogenetic reconstruction [50].

## Methods

### Simulations for training and testing CNN classifier

We used *discoal* [30] to simulate training and test datasets corresponding to each A. gambiae population under nine different selection states: neutrally evolving, completed hard sweep, completed soft sweep, partial hard sweep, partial soft sweep, and linked region for every one of the four sweep classes (Figure S1; Table S7). For the four sweep types, the target SNP was located in the exact middle position within the central, or sixth in sequence, sub-window of 11 in total; the selected SNP was placed in the middle within one of the other ten sub-windows for linked sweeps. There were 2,000 training examples per selection state (with the specific sub-window under selection, *i.e*. one of ten possible simulation classes, randomized for linked sweeps) and 1,000 test examples per class (including for each of the ten linked sweep locations), resulting in a training dataset of 18,000 simulations and a test dataset of 45,000 simulations given each demographic history, thus totaling 144,000 training and 360,000 test simulations (as well as 495,000 additional test simulations from 11 test datasets all under the BFS population history to explore demographic misspecification; see below). To conduct single-population simulations with *discoal*, we used the *stairway plot* [51] point estimates from Miles *et al*. [6] for size change parameters as well as *N*_0_ (present-day effective population size), assumed a mutation rate (*μ*) of 3.5×10^−9^ mutations per base pair per generation, and performed random draws for locus-wide mutation and recombination rates from the following distributions for each independent replicate: 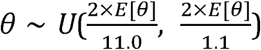, whereby *E*[*θ*] = 4*N*_0_*μL* and *L* is the length of the simulated sequence with *L* = 55,000 base pairs; *ρ* ∼ *TEXP*(2×*E*[*θ*], 6×*E*[*θ*]), where *ρ* = 4*N*_0_*rL, r* is the recombination rate per base pair, and *TEXP*(*β*, maximum value) is a truncated exponential distribution with mean *β*; *s* ∼ *U*(1.0×10^−4^, 1.0×10^−2^); end time of sweep ∼ *U*(0, 2,000) generations ago, which represents fixation for completed sweeps and the transition back to neutral evolution for partial sweeps; selected SNP allele frequency at onset of soft 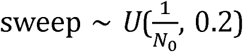; and selected SNP allele frequency at end of partial sweep ∼ *U*(0.20, 0.99).

### Constructing two-dimensional feature vector images of summary statistics

The eight training and test datasets (including an additional 11 BFS datasets for testing demographic misspecification), as well as empirical datasets, were converted into two-dimensional feature vector matrices for downstream deep learning (Figure S1); this was performed within the Python environment and required usage of the module *numpy*. Prior to this two-dimensional transformation, the simulated data were modified to better account for uncertainty within the empirical data, specifically: 1) sites that were missing any individual calls or could not be polarized against the outgroup were excluded; and 2) incorrect identification of the derived allele. For the former, each simulation randomly drew from a distribution of 1,552 masking profiles (with test simulations drawing without replacement per selection class of 1,000 simulations), which determined the exact sites to be omitted from further analysis; a masking profile consisted of the site positions within a single full 55 KB window on the *A. gambiae* genome that had absent at least one sample throughout the entirety of the Ag1000G data and/or ancestral state information, and the total set represented all 1,552 sequential, non-overlapping windows (*e.g*. 2L: 1–55,000; 2L: 55,001–110,000; 2L: 110,001–165,000; etc.) where the proportion of masked sites did not exceed 75% in any of the constituent sub-windows (*i.e*. 1,250 sites). To account for mispolarization, estimated rates were obtained from Miles *et al*. [6] and exploited via a binomial distribution to mispolarize a random subset of SNPs to the other allele per simulation.

The empirical data similarly underwent processing for compatibility with the simulated data. First, chromosomes were delineated into sequential 5 KB sub-windows (*e.g*. positions 1-5,000 formed the first sub-window, positions 5,001-10,000 formed the second sub-window, etc.), with the aforementioned masking criteria applied across sites and remaining SNPs polarized. Within each population dataset, all polymorphic positions composed of more than two alleles were further removed from analysis, such that only polarized monomorphic and biallelic sites comprising a full data matrix of no missing data were left. Sub-windows containing no SNPs or less than 25% of the original sites were subsequently discarded, and every configuration of 11 contiguous 5 KB sub-windows of those remaining formed a single full window, which would be classified into one of the nine selection states based upon its central sub-window while using spatial information from the neighboring five sub-windows on either side. To clarify, this eliminated any window that did not have a consecutive sequence of 11 sub-windows that survived data filtering, and resulted in a sliding window that progressed a single sub-window at a time, such that succeeding full windows could be overlapping by up to ten sub-windows.

Every independent simulation totaling 55 KB in length from 11 sub-windows of 5 KB, as well as empirical sequence of 11 adjacent sub-windows per population, was then transformed into 89 separate summary statistic vectors that capture aspects of population-level variation across the sampled individuals, with each vector consisting of 11 elements corresponding to the sub-windows. The first 17 summary statistics were *π* [52], *θ*_*W*_ [53], Tajima’s *D* [14], *θ*_*H*_ [15], Fay-Wu’s *H* [15], number of unique haplotypes, *H*_1_ [54], *H*_12_ [54], *H*_2_/*H*_1_ [54], *Z*_*nS*_ [55], maximum *ω* [16], *E*[*iHS*] [28], maximum *iHS* [28], proportion of outlier *iHS* values [28], variance of pairwise genotype distances [24], skewness of pairwise genotype distances [24], and kurtosis of pairwise genotype distances [24]. These were previously implemented in *S/HIC* [22] and/or *diploS/HIC* [24] except for *H*_1_ (though a multilocus genotype equivalent was used in *diploS/HIC*) and the *iHS*-based statistics. We employed the Python package scikit-allel to calculate *π, θ*_*W*_, Tajima’s *D, θ*_*H*_, Fay-Wu’s *H*, number of unique haplotypes, *H*_1_, *H*_12_, *H*_2_/*H*_1_, and the statistics related to *iHS*. Values for *iHS* were standardized within 50 derived allele frequency bins, following mispolarization in the case of simulated data. Outlier *iHS* values were defined as within either 2.0% tail of the distribution obtained from simulations of neutral evolution under the appropriate demographic history.

The remaining 72 summary statistics were distribution summaries of SAFE and its various components [29], specifically: haplotype allele frequency (HAF), which is the sum of derived allele counts across all the derived alleles present within a sequence; unique HAF score (*i.e*. each unique HAF value is counted only once, even if representing multiple individuals); *φ*, which is the sum of HAF scores for sequences harboring the derived allele, divided by the total sum of HAF scores across all sequences; κ, which is the proportion of distinct HAF scores that carry the derived allele; derived allele frequency; and the SAFE score itself, which is the difference between *φ* and κ normalized against the derived allele frequency. Notably, HAF is calculated per sequence, whereas *φ*, κ, derived allele frequency, and SAFE are calculated per polymorphism. The following distribution summaries were utilized to construct individual values spanning a sub-window: mean, median, mode; 2.5%, 25%, 75%, and 97.5% quartiles; maximum, variance, standard deviation, skewness, and kurtosis. Importantly, each summary statistic vector was normalized, in the same manner as the preceding versions to *partialS/HIC* [22,24], to capture signal solely from the relative spatial distribution of the summary statistics across the 11 sub-windows, rather than allowing influence from absolute values. Subsequently, the 89 vectors were vertically collated to form a two-dimensional matrix that could then be exploited for image processing. The arrangement of these vectors were such that the 11 columns corresponded to the series of sub-windows from left to right, and the 89 rows of summary statistics were in the order presented here (with the distribution summaries iterating first for every SAFE component, *e.g*. row 52: skewness of *φ* values; row 53: kurtosis of *φ* values; row 54: *E*[κ]). Importantly, column and row order affects deep learning optimization, which may have consequences on overall efficacy, due to the convolutional and pooling windows employed by the CNN architecture, hence related summary statistics were grouped together (*e.g*. alternative distribution summaries of a SNP-based statistic, various SAFE derivatives). Heatmap images, based on median values per statistic and sub-window, were generated in R for the neutral case and each of the four sweep states under every population history from the training simulations.

### Training and testing CNNs for deep learning implementation

The architecture of our CNN was composed of the following sequential layers: 1) 2D convolutional layer with 256 filters using 3×6 windows and “same” padding; 2) 2D max pooling layer given a 3×3 window; 3) a second 2D convolutional layer of 256 filters based also on 3×3 windows, “same” padding, and ReLU activation; 4) a second 2D max pooling layer with a 3×1 window; 5) dropout layer with p=0.25; 6) flattening layer; 7) fully-connected layer with ReLU activation to 512 responses; 8) a second dropout layer with p=0.50; 9) a second fully-connected layer with ReLU activation to 128 elements; 10) another dropout layer with p=0.50; and 11) softmax activation layer to 9 states (Figure 1). This architecture was trained using the Python module *Keras* [56] given the *Adam* optimizer [57], with 20 epochs, batch size of 32 simulations per step within an epoch, and 10% of the training data (*e.g*. 1800 simulations from the total nine-state training dataset) randomly removed as a validation set during optimization (Figure S1). Training was performed for every population demography under three experimental settings: 1) given the full set of training data distributed across nine selection states; 2) exploiting a subset of the training data from only five of the selection states, specifically those involving neutral regions or completed sweeps; 3) deploying the entire training data, but with binary classification between selective sweeps in the focal sub-window and all unselected classes (*i.e*. neutral class together with every linked class). The three training regimes were then applied to the test dataset that corresponded in underlying simulated history, resulting in predictions of selection state given the default *Keras* threshold parameters for the first two training schemes, whereas the softmax probability of a focal sweep was instead exploited under training on binary classification. In the former case, individual inferences per test simulation were collated and compared against true values to produce overall accuracy measures as well as confusion matrix heatmaps to assess misclassification bias for each of the 45 simulated scenarios (*i.e*. 11 sub-window sweep locations for each of the four sweep types, plus neutrality).

Moreover, to explore the effect of demographic misspecification, we conducted a single additional test under the first experimental set-up whereby the CNN trained on the GAS simulations was applied to the CMS test simulations. Furthermore, we engaged in two more misspecification experiments that involved simulating, for the first experiment, five testing datasets that changed the underlying demographic model, and for the second experiment, six testing datasets that differed in fixed *θ* value. In the five test sets that altered the demographic history, we used an instantaneous contraction model that experienced a population crash at 0.0001 time units with an intensity of, respectively across the five datasets: 20×, 40×, 60×, 80×, 100×. Priors for *θ, ρ, s*, end time of sweep, selected SNP allele frequency at onset of soft sweep, and selected SNP allele frequency at end of partial sweep remained the same as for the initial BFS simulations. In the six test sets that varied *θ*, we employed the same parameterization from the initial BFS simulations with the exception that *θ* was set to, respectively across the six datasets: 1,000; 5,000; 10,000; 15,000; 20,000; 25,000. Our motivation for these intervals was to exceed the bounds of the original prior distribution, *θ* ∼ *U*(1,750.204699, 17,502.046985). After simulation, we leveraged the nine-state CNN trained under the BFS specifications against each of the 11 testing datasets to obtain overall accuracies for comparison among these test datasets and to the original test dataset whereby the BFS history was correctly modeled.

Regarding the binary classification experiment, the Python module *sklearn* [58] was used for building the ROC curves to evaluate accuracy and sensitivity. Additionally, *sklearn* was utilized for conducting PCA on the training simulations, on either the focal sub-window or across the entire full-scale genomic window respectively, with subsequent projection of the testing simulations into independent sets of PC1 and PC2 values, to obtain a total of four individual ROC curves. To calculate the Composite of Multiple Signals [31], training data were separated into the two categories of selection versus neutrality (*i.e*. 4,000 simulations from four selection states versus 41,000 simulations from 41 neutral or linked classes), which were respectively converted into a histogram of 200 bins for every summary statistic. Assuming the Composite of Multiple Signals for the central sub-window only, this resulted in a total of 178 distributions given the 89 statistics for both selection and neutral; for all sub-windows, this was instead a total of 1,958 distributions due to the 11 total sub-windows of statistics. These distributions were then employed to calculate probabilities both under selection and neutrality for the corresponding testing data summary statistics; for bins with a zero value, 0.1 of the smallest possible increment (*i.e*.selection: 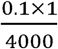; neutrality: 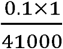) was assigned as the probability. To determine the prior for each probability, represented in the original Composite of Multiple Signals equation by *π*, we utilized the proportional composition of the training simulations (*i.e*. selection: 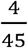; neutrality: 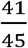). Finally, probabilities were natural log transformed and summed among summary statistics to compute the Composite of Multiple Signals.

### Detecting selective sweeps for *A. gambiae* population datasets

To scan the genome for signatures of selective sweeps, the nine-state trained CNNs were applied to the eight empirical mosquito datasets, with the underlying simulated demography matched to the sampled population (Figure S1). Calls were corrected for false discovery by exploiting the accuracy and error rates for neutral regions from the nine-state simulation experiment, such that the amount of neutral calls was assumed to be underestimated while the amount of calls for the remaining eight selection states were assumed to be inflated. Subsequently, we produced sets of 10,000 randomly permuted calls across the genome to derive null expectations of sweep enrichment, following Schrider and Kern [32]. Using the gene annotation file “Anopheles-gambiae-PEST_BASEFEATURES_AgamP4.7.gff3.gz” from VectorBase, we exploited these permuted datasets to assess statistically significant enrichment within certain DNA regions, groupings of known IR genes (N. Harding, *pers. comm*.), and all basic GO term definitions from http://www.geneontology.org (last accessed February 18, 2015). The DNA regions of interest included gene, mRNA, exon, CDS, five-prime UTR, and three-prime UTR; IR genes were assigned to four functional categories: metabolism, target sites, behavior, and cuticular. To determine significant enrichment, the number of inferred calls for a particular DNA region or IR gene category had to have a *p*-value < 0.05 based on the respective distribution of 10,000 permutations; for the GO terms, we deployed a corrected *q*-value < 0.05 due to concerns of false discovery stemming from the large number of terms tested for enrichment.

## Acknowledgments

We thank Jeff Adrion for comments on the manuscript. This work was supported by NIH award no. R01GM117241 to ADK and NIH award no. K99HG008696 to DRS.

## Figure Captions

**Figure S1. Flowchart of *partialS/HIC* methodological procedure for training, testing, and empirical application given eight *Anopheles* mosquito population datasets.** Major steps highlighted in green, specifically data simulation, training and testing of CNN classifier, and genome-wide inference of selection state per focal sub-window within a full window of 11 sequential sub-windows. Important data processing details that define genomic sub-windows as well as account for uncertainties within the empirical data from missing sites and polarization are also included. However, we omit our various misclassification experiments, which are ancillary to our primary demonstration of *partialS/HIC* to detect selective sweeps from population genomic datasets while obtaining baseline measures of accuracy. For organizational and conceptual purposes, our protocol is presented here mostly contained within two columns, representing simulated versus empirical data respectively, but the actual order of events oscillates between these two sides of the diagram. In fact, many of the tasks could be accomplished in parallel, though certain actions must occur sequentially. For example, training and testing data were generated after demographic inference in order to inform parameterization of the simulations. Moreover, we filtered the empirical data of missing calls and missing polarization information prior to constructing masking profiles for application to the simulated data, which invoked an intermediate step hence our usage of a dotted arrow. Additionally, the CNN first had to be optimized from the training simulations before conducting empirical analysis.

**Figure S2. Feature vector heatmaps for neutral and four sweep states under each population history.** Heatmaps derived from median values of a single training dataset, each assuming a different population history. Across populations, feature vectors demonstrate distinctive characteristics per classification state, especially in the focal sub-window and for the neutral state in comparison to the four sweep states.

**Figure S3. Training and validation loss over samples and epochs during *partialS/HIC* CNN optimization under each population history.** Black circles represent loss function outputs from training for every 32 samples out of a total 16,200 samples per each of 20 epochs. Green asterisks indicate loss function outputs from the validation set at the conclusion of epochs. The degree to which training loss is less than validation loss likely reflects the magnitude of overfitting the CNN to the training data.

**Figure S4. Confusion matrix heatmaps of *partialS/HIC* simulation experiment under each population history.** Extension of Figure 2 for all eight population histories. Importantly, the patterns highlighted in Figure 2, which displays the confusion matrix given the BFS population history and is reproduced identically here, are largely present throughout the other confusion matrices.

**Figure S5. Confusion matrix heatmap of *partialS/HIC* demographic misspecification test.** Training on simulations given the GAS population history while testing on simulations given the CMS population history. Importantly, the patterns highlighted in Figures 2 and S4 are largely retained here, though there is a modest decrease in soft sweep discovery for both completed and partial.

**Figure S6. Overall *partialS/HIC* accuracies between testing datasets generated with varying demographic misspecification given training on the BFS population history.** The CNN trained on the simulations assuming the BFS population history was separately applied against five testing datasets that were produced under a simple two-epoch instantaneous population crash model, with each dataset differing only in relative magnitude of the size change. The overall accuracy from the original testing dataset whereby the demography was accurately specified on the BFS population history is marked by the green dotted line at 67.76%. The final population size ranges from 5% (*i.e*. 20×) to 1% (*i.e*. 100×) of the original size, with increased accuracy for more severe contractions (*i.e*. 45.98% accuracy for reduction to 5.00% of original size; 55.27% for reduction to 2.50%; 59.46% for reduction to 1.67%; 61.99% for reduction to 1.25%; and 63.50% for reduction to 1.00%). Notably, the greater error at lower magnitudes could be largely due to the decrease in simulated polymorphisms (Figure S7).

**Figure S7. Overall *partialS/HIC* accuracies between testing datasets generated with varying *θ* values given training on the BFS population history.** The CNN trained on the simulations assuming the BFS population history was separately applied against six testing datasets that were produced under identical conditions as the training data except differing in fixed *θ* values. The overall accuracy from the original testing dataset whereby *θ* was drawn from a prior distribution is marked by the green dotted line at 67.76%. Across values of 1,000, 5,000, 10,000, 15,000, 20,000, and 25,000, which collectively is inclusive of and exceeds the original prior distribution (*θ* ∼ *U*(1,750.204699, 17,502.046985)), there is comparable accuracy to the original (50.99%, 67.78%, 70.36%, 70.74%, 70.72%, and 70.74% respectively). In fact, accuracy exceeded the original for all tested *θ* values except *θ* = 1,000.

**Figure S8. Confusion matrix heatmaps of simulation experiment with partial sweeps ignored during training under each population history.** Extension of Figure 3 for all eight population histories. Importantly, the patterns highlighted in Figure 3, which displays the confusion matrix given the BFS population history and is reproduced identically here, are largely present throughout the other confusion matrices.

**Figure S9. ROC curves for binary classification of selective sweeps, including partial sweeps, versus regions neutrally evolving or under linked selection for each population history.** Extension of Figure 4 for all eight population histories. Importantly, the patterns highlighted in Figure 4, which displays the ROC curves given the BFS population history and is reproduced identically here, are largely present throughout the other plots of ROC curves.

**Figure S10. Enrichments of *partialS/HIC* empirical selective sweep calls within DNA regions for each empirical mosquito population dataset.** Extension of Figure 6 for all eight empirical population datasets. Importantly, the patterns highlighted in Figure 6, which displays the dot chart for the BFS permutation test and is reproduced here with a wider X-axis to match the other plots, are largely present throughout the other dot charts, though partial sweep enrichment is highly variable.

**Figure S11. Enrichments of *partialS/HIC* empirical selective sweep calls within groupings of insecticide resistance genes for each empirical mosquito population dataset.** Extension of Figure 7 for all eight empirical population datasets. Importantly, as in Figure 7, which displays the dot chart for the BFS permutation test and is reproduced identically here, there is evidence for several mechanisms through which insecticide resistance evolved, both in sweep types and categories of genes under selection as well as both within individual populations and between different populations.

