## Supplemental Figures for "Discovery of ongoing selective sweeps within *Anopheles* mosquito populations using deep learning"

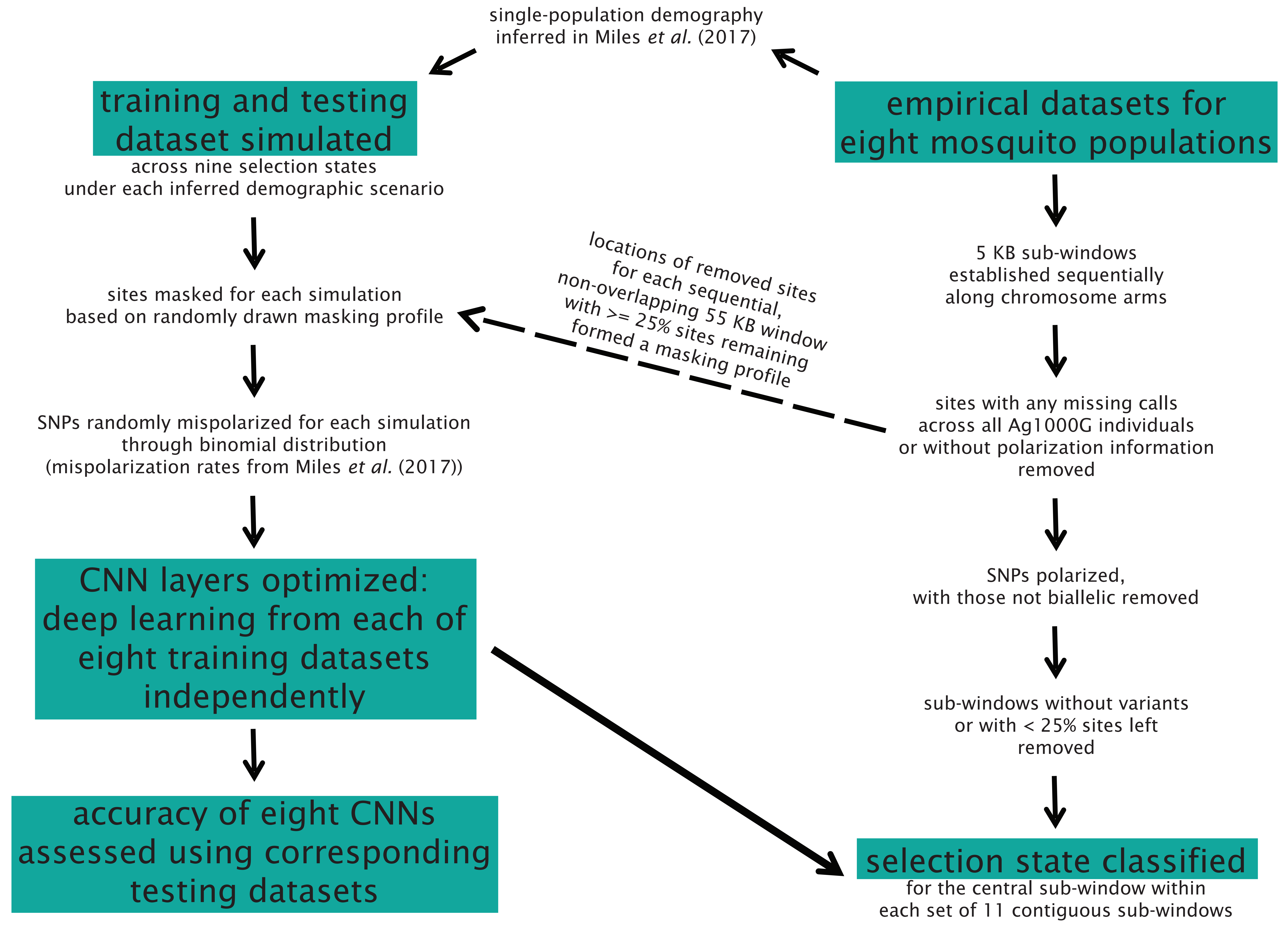
single-population demography
inferred in Miles *et al.* (2017)
training and testing
dataset simulated
across nine selection states
under each inferred demographic scenario
sites masked for each simulation
based on randomly drawn masking profile
SNPs randomly mispolarized for each simulation
through binomial distribution
(mispolarization rates from Miles *et al.* (2017))
CNN layers optimized:
deep learning from each of
eight training datasets
independently
accuracy of eight CNNs
assessed using corresponding
testing datasets
empirical datasets for
eight mosquito populations
5 KB sub-windows
established sequentially
along chromosome arms
locations of removed sites
for each sequential,
non-overlapping 55 KB window
with >= 25% sites remaining
formed a masking profile
sites with any missing calls
across all Ag1000G individuals
or without polarization information
removed
SNPs polarized,
with those not biallelic removed
sub-windows without variants
or with < 25% sites left
removed
selection state classified
for the central sub-window within
each set of 11 contiguous sub-windows

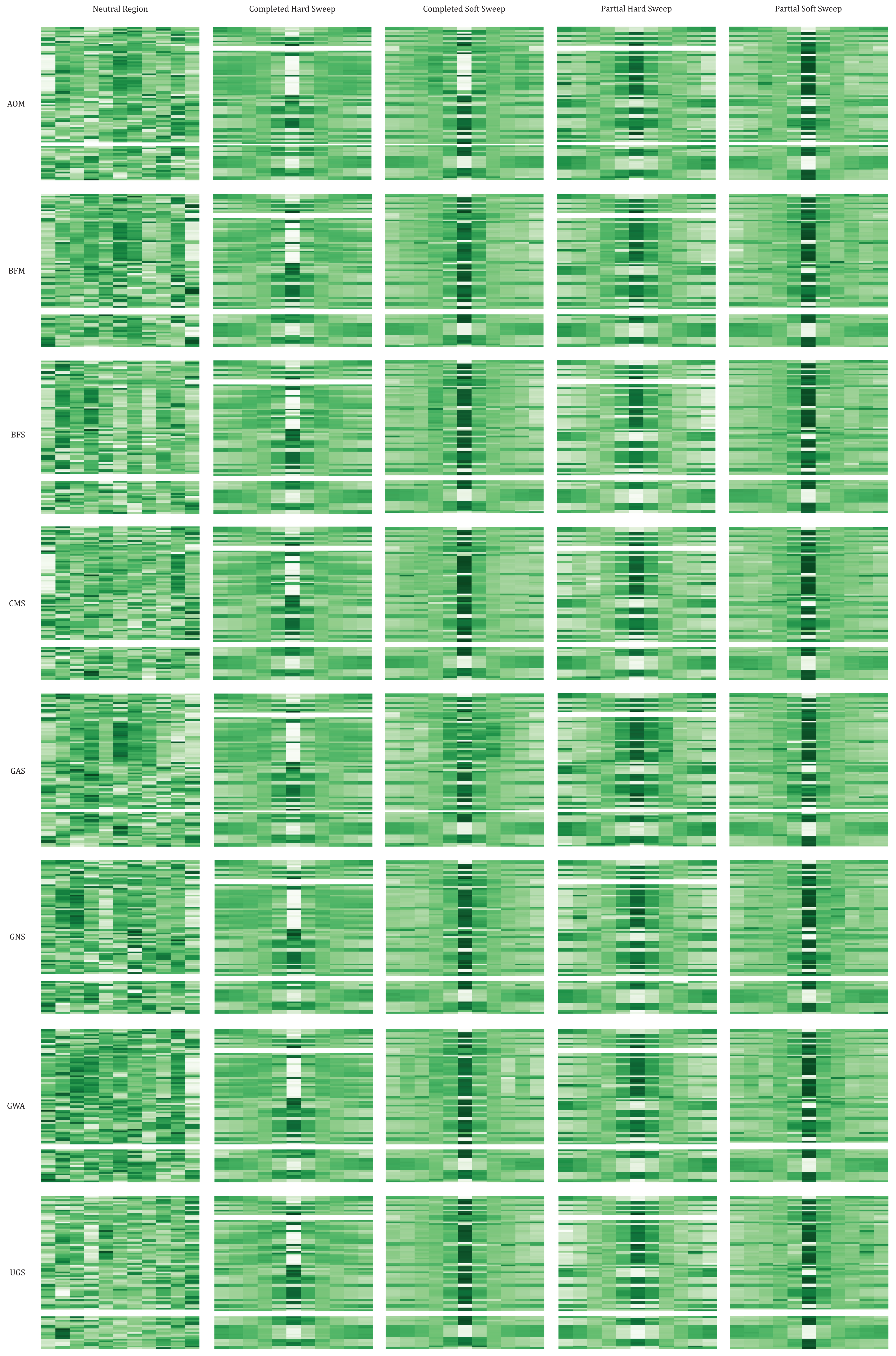
Neutral Region
Completed Hard Sweep
Completed Soft Sweep
Partial Hard Sweep
Partial Soft Sweep
AOM
BFM
BFS
CMS
GAS
GNS
GWA
UGS

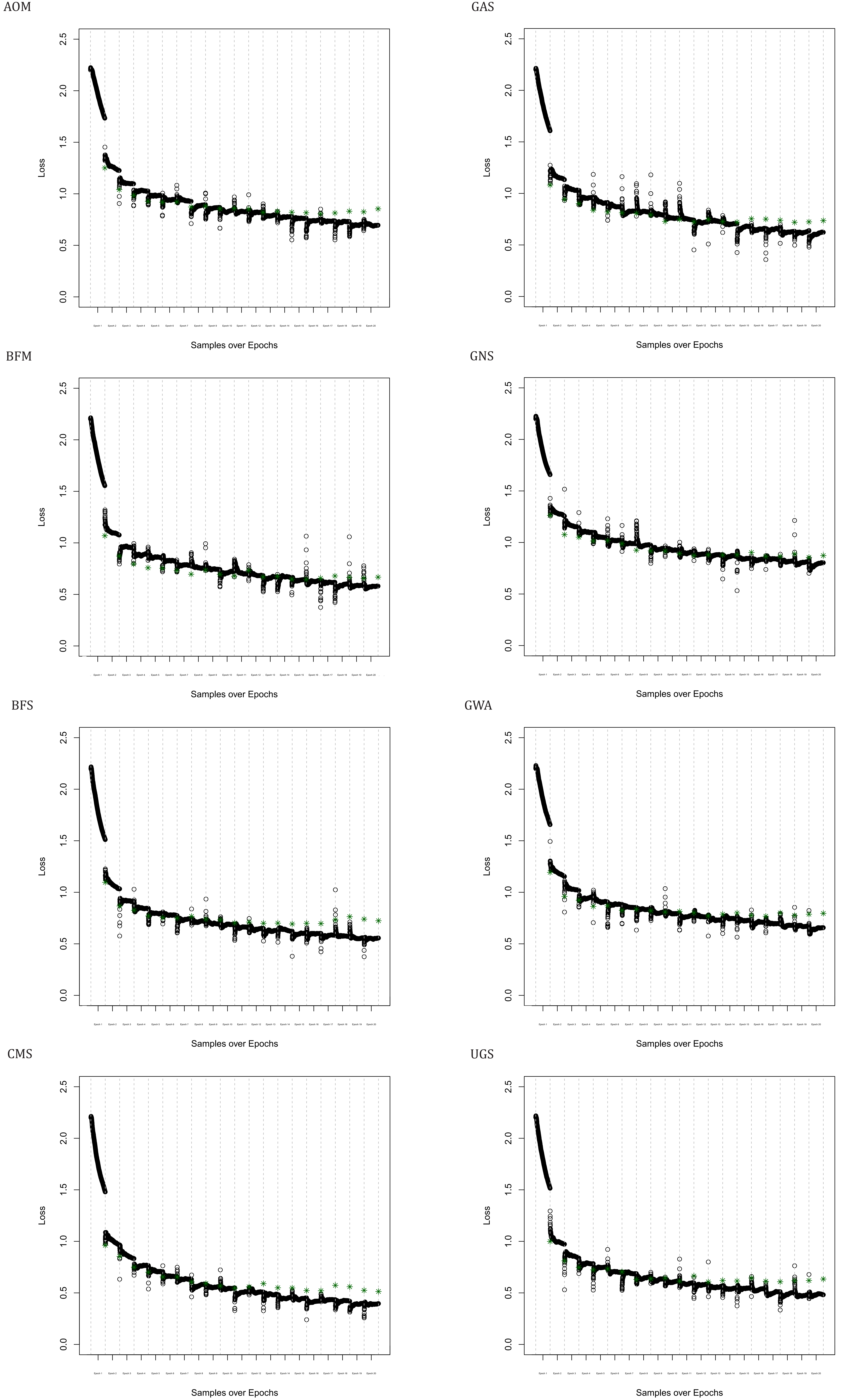

AOM
GAS
BFM
GNS
BFS
GWA
CMS
UGS
Loss
Samples over Epochs

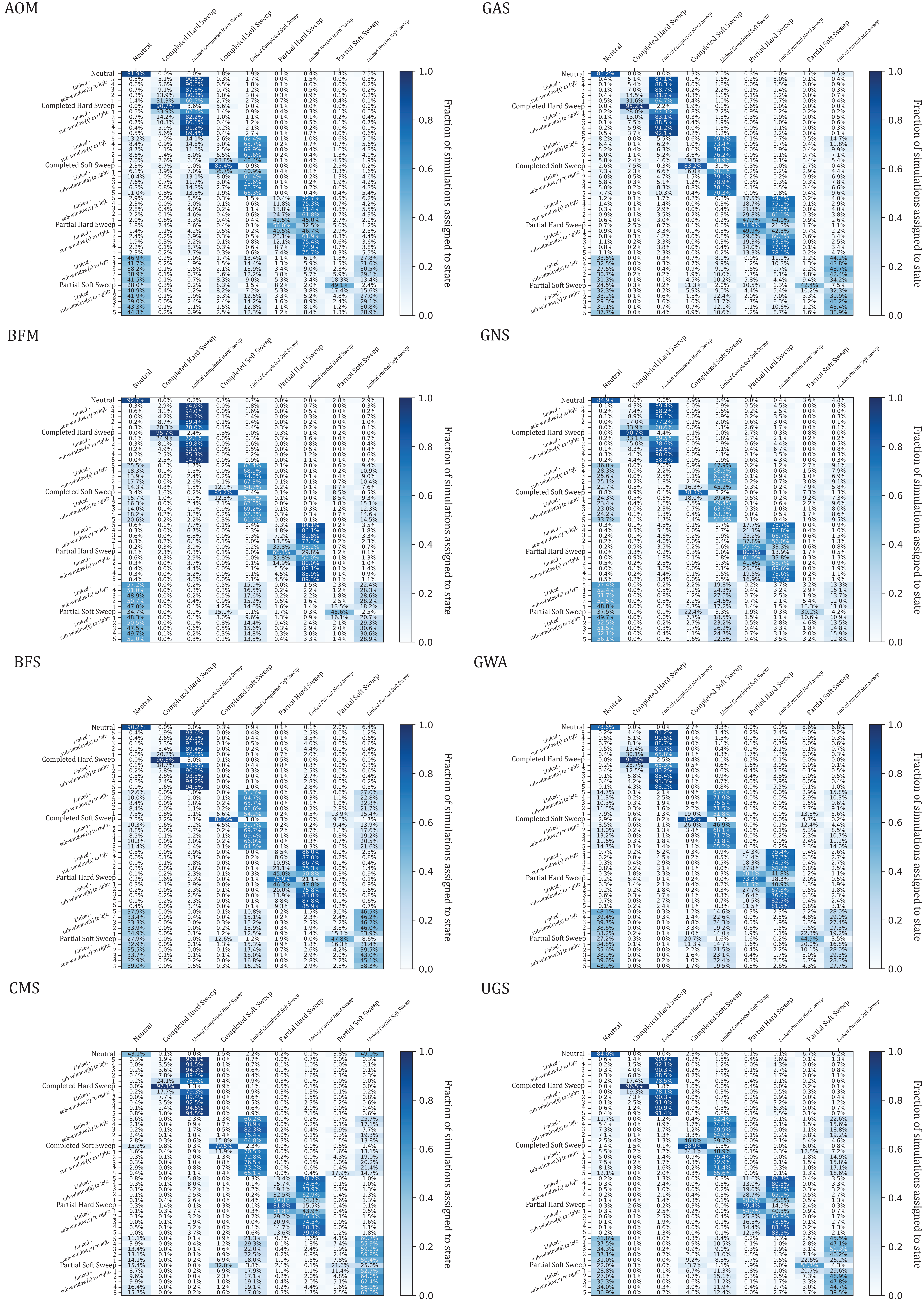

AOM
GAS
BFM
GNS
BFS
GWA
CMS
UGS
Neutral
Completed Hard Sweep
Linked Completed Hard Sweep
Completed Soft Sweep
Linked Completed Soft Sweep
Partial Hard Sweep
Linked Partial Hard Sweep
Partial Soft Sweep
Linked Partial Soft Sweep
Linked - sub-window(s) to left:
Linked - sub-window(s) to right:
Fraction of simulations assigned to state
1.0
0.8
0.6
0.4
0.2
0.0

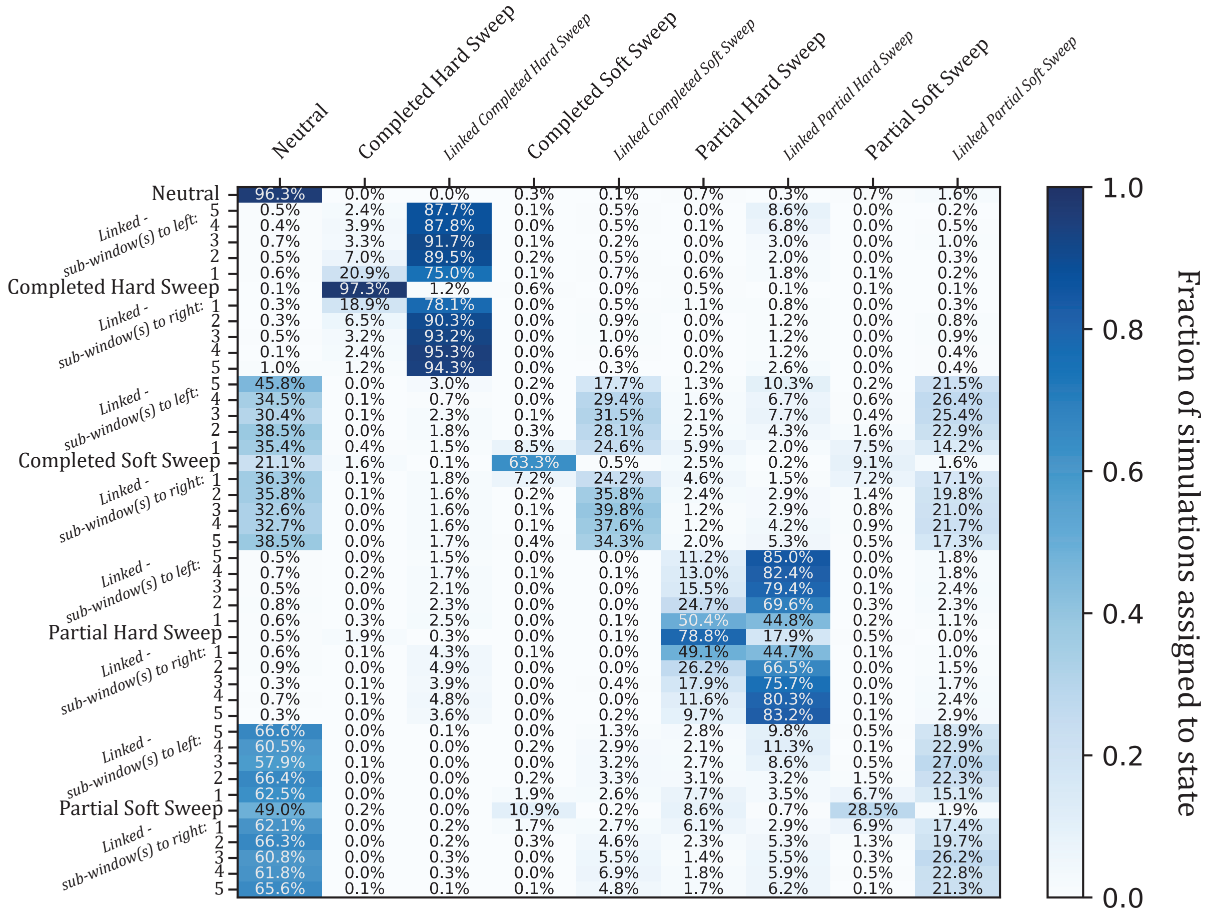

Neutral
Completed Hard Sweep
Linked Completed Hard Sweep
Completed Soft Sweep
Linked Completed Soft Sweep
Partial Hard Sweep
Linked Partial Hard Sweep
Partial Soft Sweep
Linked Partial Soft Sweep
Neutral 96.3% 0.0% 0.0% 0.3% 0.1% 0.7% 0.3% 0.7% 1.6%
Linked - sub-window(s) to left:
5 0.5% 2.4% 87.7% 0.1% 0.5% 0.0% 8.6% 0.0% 0.2%
4 0.4% 3.9% 87.8% 0.0% 0.5% 0.1% 6.8% 0.0% 0.5%
3 0.7% 3.3% 91.7% 0.1% 0.2% 0.0% 3.0% 0.0% 1.0%
2 0.5% 7.0% 89.5% 0.2% 0.5% 0.0% 2.0% 0.0% 0.3%
1 0.6% 20.9% 75.0% 0.1% 0.7% 0.6% 1.8% 0.1% 0.2%
Completed Hard Sweep 0.1% 97.3% 1.2% 0.6% 0.0% 0.5% 0.1% 0.1% 0.1%
Linked - sub-window(s) to right:
1 0.3% 18.9% 78.1% 0.0% 0.5% 1.1% 0.8% 0.0% 0.3%
2 0.3% 6.5% 90.3% 0.0% 0.9% 0.0% 1.2% 0.0% 0.8%
3 0.5% 3.2% 93.2% 0.0% 1.0% 0.0% 1.2% 0.0% 0.9%
4 0.1% 2.4% 95.3% 0.0% 0.6% 0.0% 1.2% 0.0% 0.4%
5 1.0% 1.2% 94.3% 0.0% 0.3% 0.2% 2.6% 0.0% 0.4%
Linked - sub-window(s) to left:
5 45.8% 0.0% 3.0% 0.2% 17.7% 1.3% 10.3% 0.2% 21.5%
4 34.5% 0.1% 0.7% 0.0% 29.4% 1.6% 6.7% 0.6% 26.4%
3 30.4% 0.1% 2.3% 0.1% 31.5% 2.1% 7.7% 0.4% 25.4%
2 38.5% 0.0% 1.8% 0.3% 28.1% 2.5% 4.3% 1.6% 22.9%
1 35.4% 0.4% 1.5% 8.5% 24.6% 5.9% 2.0% 7.5% 14.2%
Completed Soft Sweep 21.1% 1.6% 0.1% 63.3% 0.5% 2.5% 0.2% 9.1% 1.6%
Linked - sub-window(s) to right:
1 36.3% 0.1% 1.8% 7.2% 24.2% 4.6% 1.5% 7.2% 17.1%
2 35.8% 0.1% 1.6% 0.2% 35.8% 2.4% 2.9% 1.4% 19.8%
3 32.6% 0.0% 1.6% 0.1% 39.8% 1.2% 2.9% 0.8% 21.0%
4 32.7% 0.0% 1.6% 0.1% 37.6% 1.2% 4.2% 0.9% 21.7%
5 38.5% 0.0% 1.7% 0.4% 34.3% 2.0% 5.3% 0.5% 17.3%
Linked - sub-window(s) to left:
5 0.5% 0.0% 1.5% 0.0% 0.0% 11.2% 85.0% 0.0% 1.8%
4 0.7% 0.2% 1.7% 0.1% 0.1% 13.0% 82.4% 0.0% 1.8%
3 0.5% 0.0% 2.1% 0.0% 0.0% 15.5% 79.4% 0.1% 2.4%
2 0.8% 0.0% 2.3% 0.0% 0.0% 24.7% 69.6% 0.3% 2.3%
1 0.6% 0.3% 2.5% 0.0% 0.1% 50.4% 44.8% 0.2% 1.1%
Partial Hard Sweep 0.5% 1.9% 0.3% 0.0% 0.1% 78.8% 17.9% 0.5% 0.0%
Linked - sub-window(s) to right:
1 0.6% 0.1% 4.3% 0.1% 0.0% 49.1% 44.7% 0.1% 1.0%
2 0.9% 0.0% 4.9% 0.0% 0.0% 26.2% 66.5% 0.0% 1.5%
3 0.3% 0.1% 3.9% 0.0% 0.4% 17.9% 75.7% 0.0% 1.7%
4 0.7% 0.1% 4.8% 0.0% 0.0% 11.6% 80.3% 0.1% 2.4%
5 0.3% 0.0% 3.6% 0.0% 0.2% 9.7% 83.2% 0.1% 2.9%
Linked - sub-window(s) to left:
5 66.6% 0.0% 0.1% 0.0% 1.3% 2.8% 9.8% 0.5% 18.9%
4 60.5% 0.0% 0.0% 0.2% 2.9% 2.1% 11.3% 0.1% 22.9%
3 57.9% 0.1% 0.0% 0.0% 3.2% 2.7% 8.6% 0.5% 27.0%
2 66.4% 0.0% 0.0% 0.2% 3.3% 3.1% 3.2% 1.5% 22.3%
1 62.5% 0.0% 0.0% 1.9% 2.6% 7.7% 3.5% 6.7% 15.1%
Partial Soft Sweep 49.0% 0.2% 0.0% 10.9% 0.2% 8.6% 0.7% 28.5% 1.9%
Linked - sub-window(s) to right:
1 62.1% 0.0% 0.2% 1.7% 2.7% 6.1% 2.9% 6.9% 17.4%
2 66.3% 0.0% 0.2% 0.3% 4.6% 2.3% 5.3% 1.3% 19.7%
3 60.8% 0.0% 0.3% 0.0% 5.5% 1.4% 5.5% 0.3% 26.2%
4 61.8% 0.0% 0.3% 0.0% 6.9% 1.8% 5.9% 0.5% 22.8%
5 65.6% 0.1% 0.1% 0.1% 4.8% 1.7% 6.2% 0.1% 21.3%
1.0
0.8
0.6
0.4
0.2
0.0
Fraction of simulations assigned to state

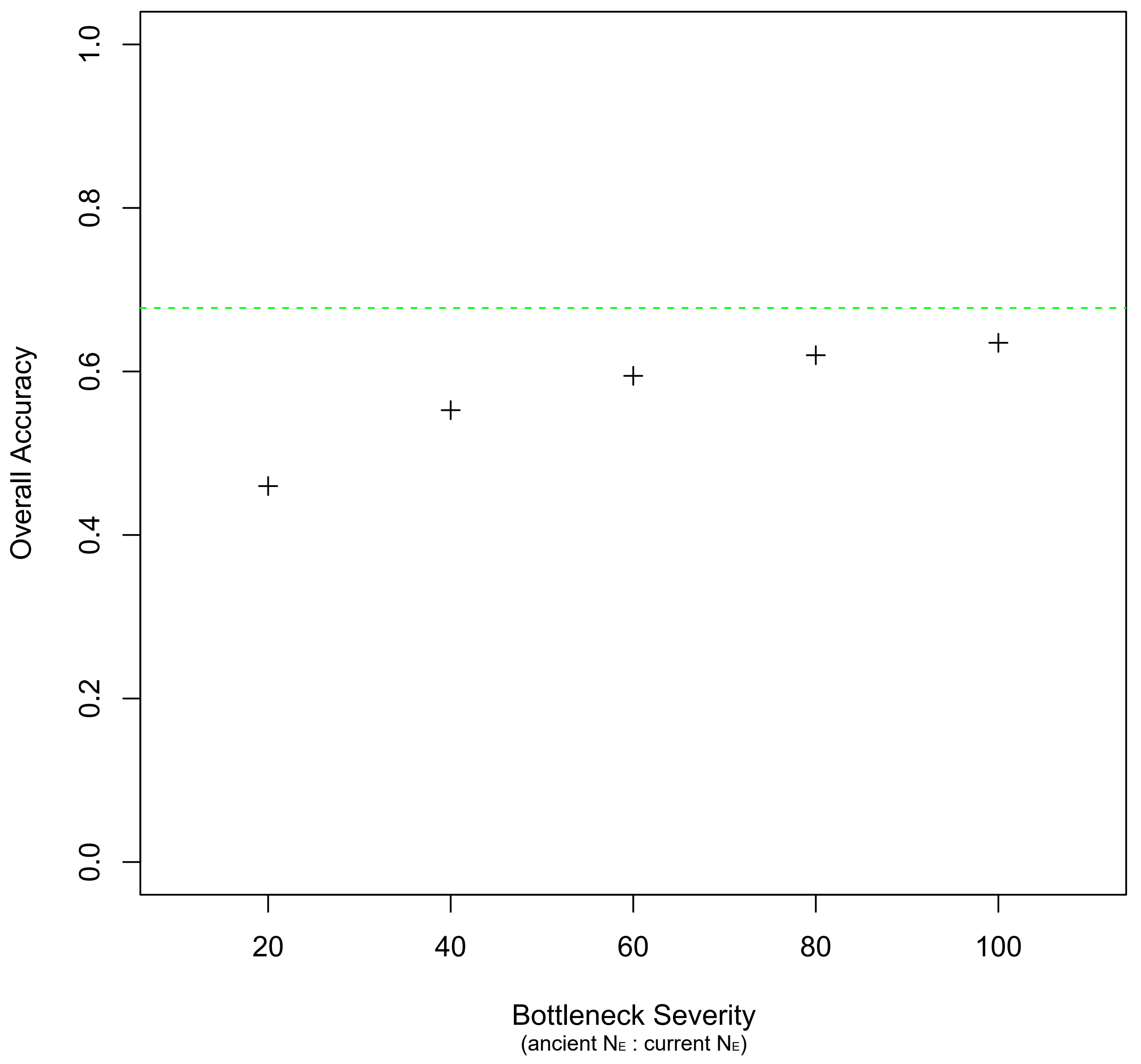

Overall Accuracy
1.0
0.8
0.6
0.4
0.2
0.0
20
40
60
80
100
Bottleneck Severity
(ancient NE : current NE)

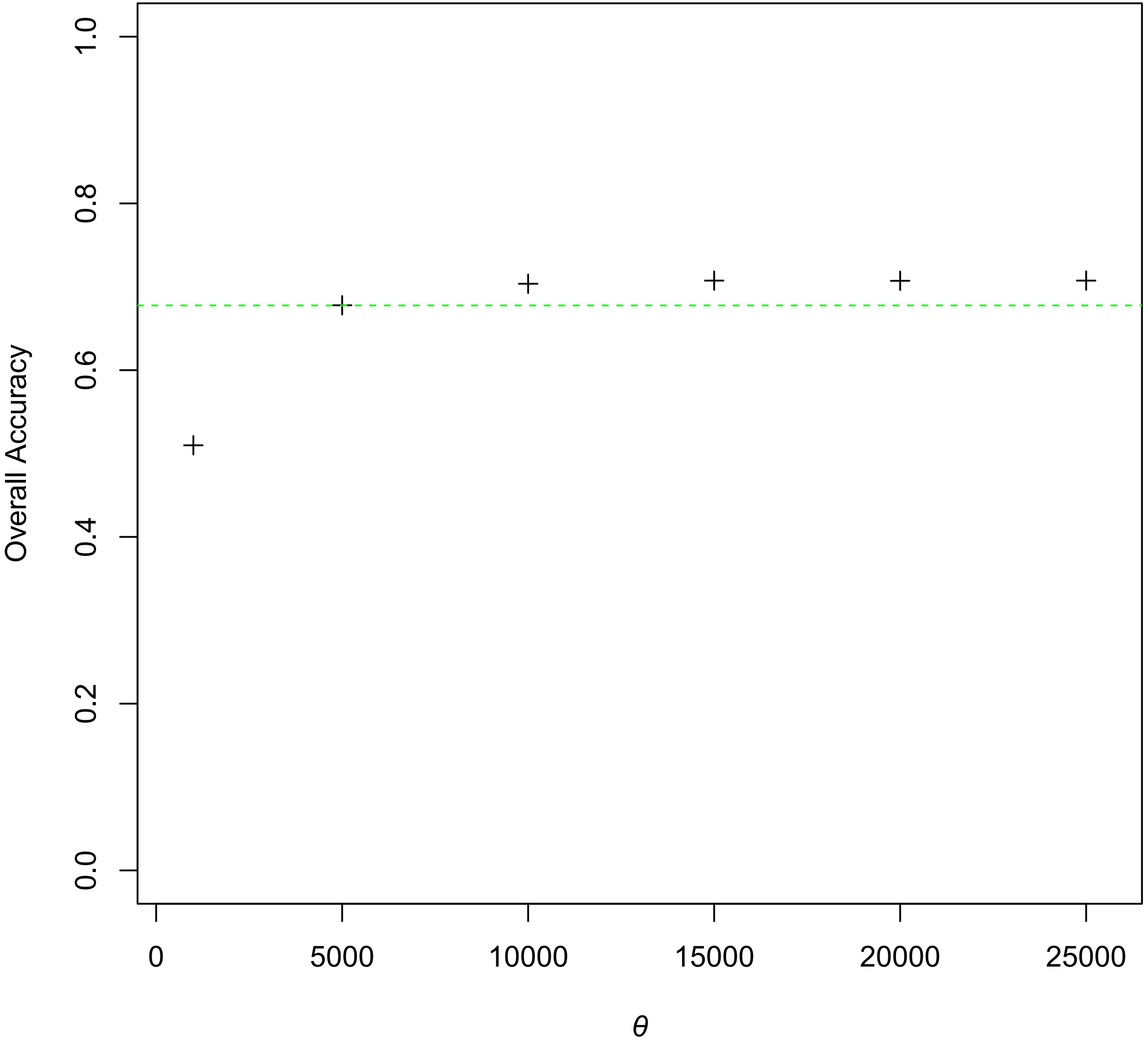

Overall Accuracy
0.0
0.2
0.4
0.6
0.8
1.0
0
5000
10000
15000
20000
25000
θ

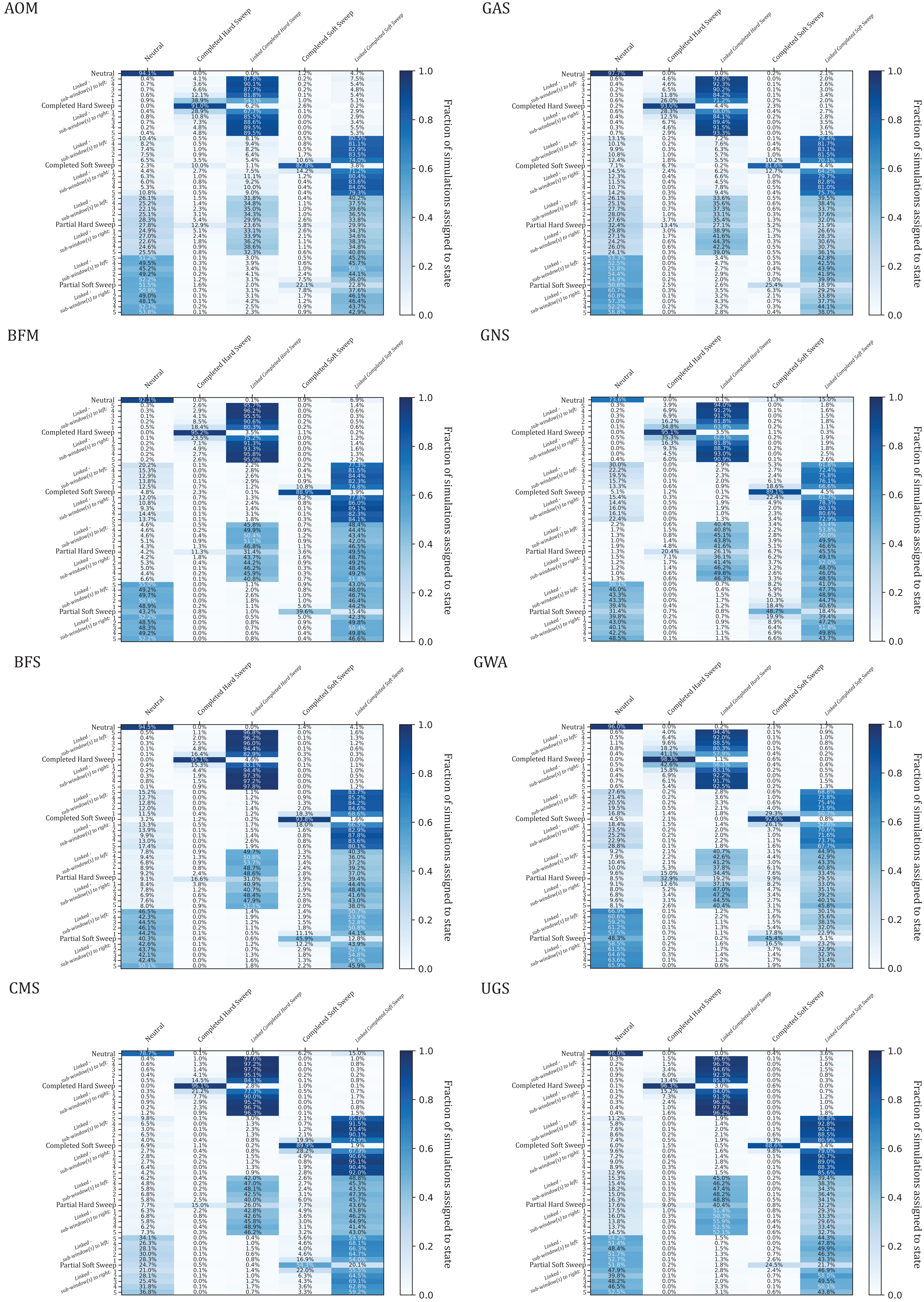
AOM
GAS
BFM
GNS
BFS
GWA
CMS
UGS
Neutral
Completed Hard Sweep
Linked Completed Hard Sweep
Completed Soft Sweep
Linked Completed Soft Sweep
Partial Hard Sweep
Partial Soft Sweep
Linked - sub-window(s) to left:
Linked - sub-window(s) to right:
Fraction of simulations assigned to state
0.0
0.2
0.4
0.6
0.8
1.0

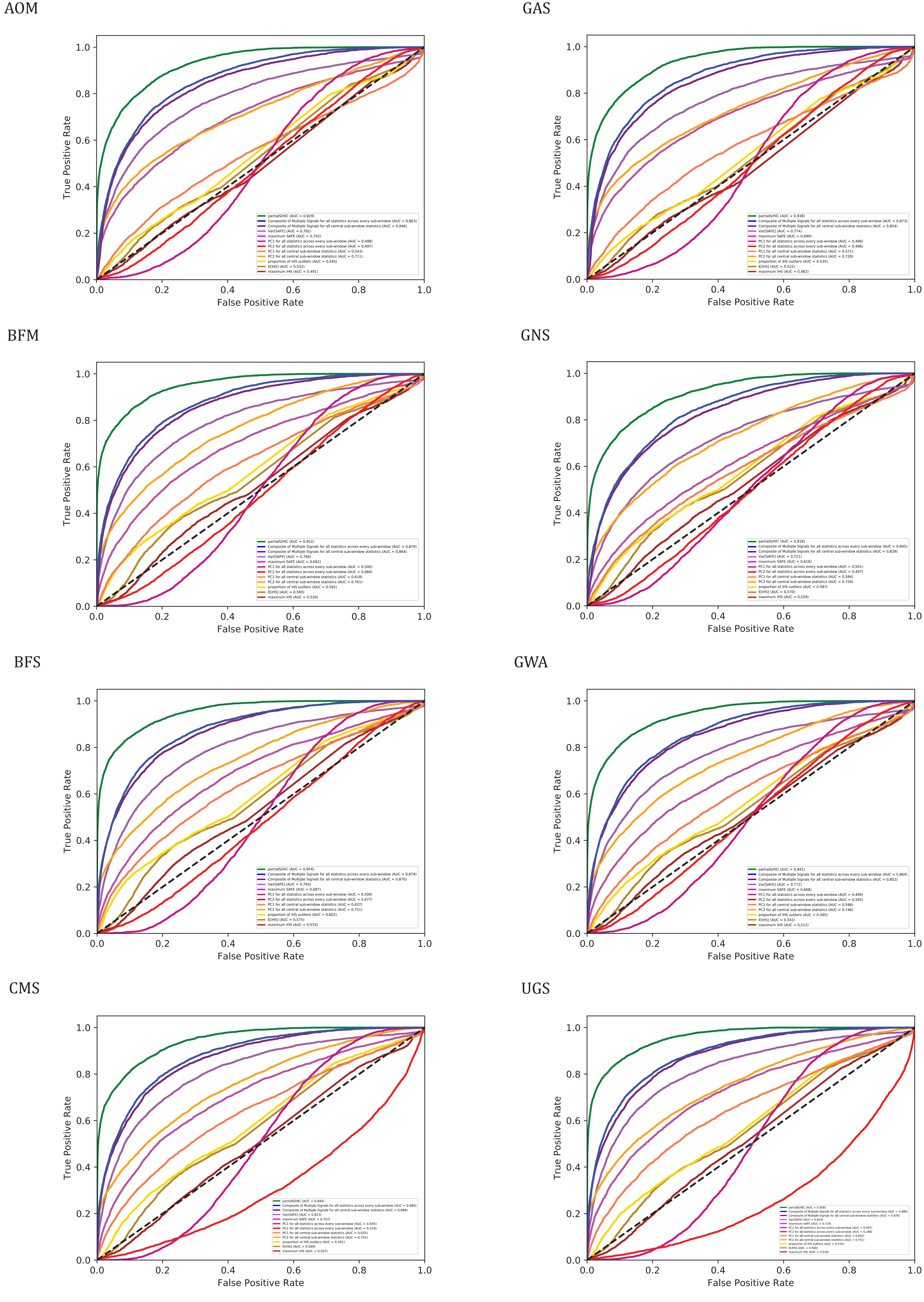

AOM
GAS
BFM
GNS
BFS
GWA
CMS
UGS
True Positive Rate
False Positive Rate

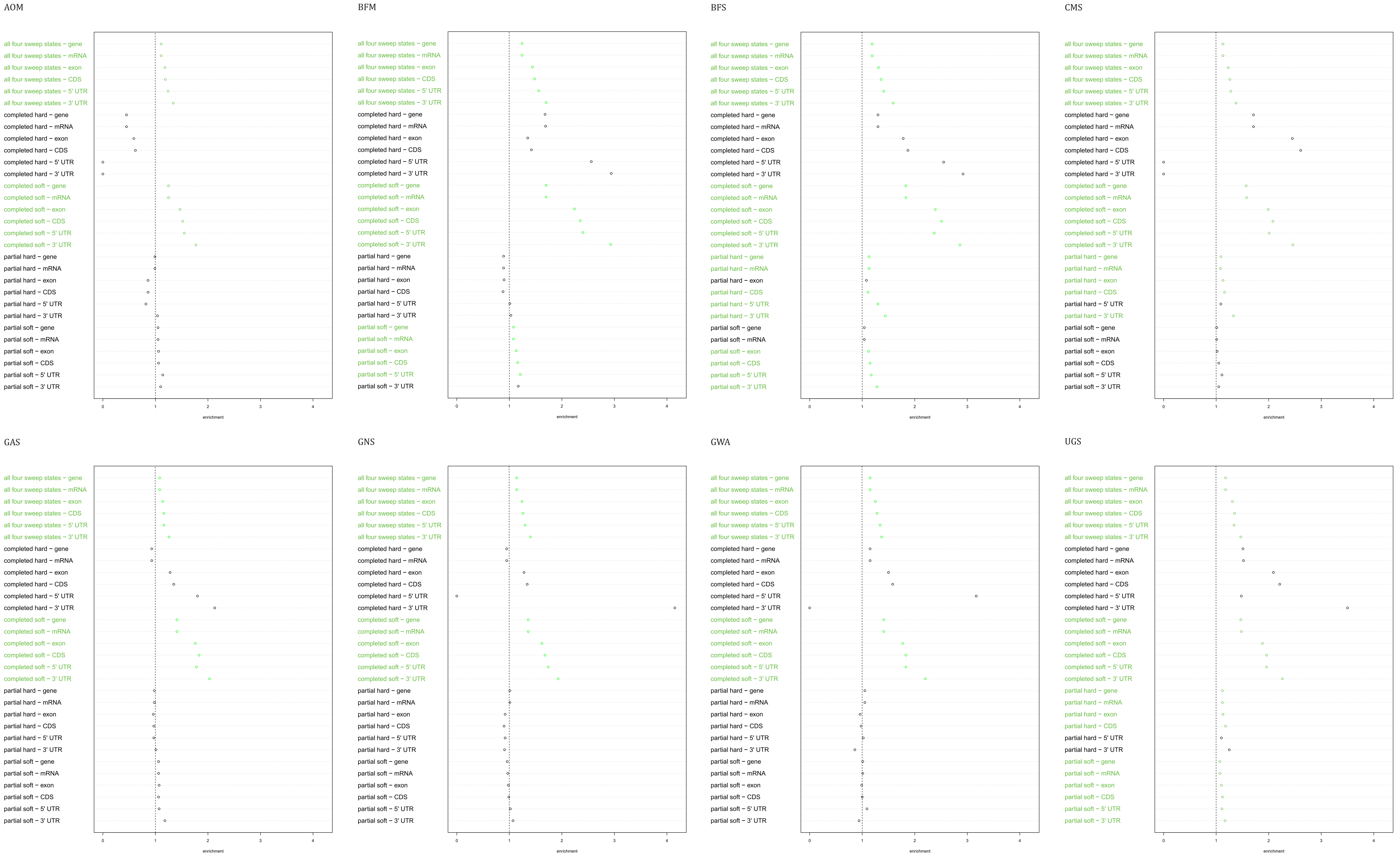

AOM
BFM
BFS
CMS
GAS
GNS
GWA
UGS
all four sweep states − gene
all four sweep states − mRNA
all four sweep states − exon
all four sweep states − CDS
all four sweep states − 5' UTR
all four sweep states − 3' UTR
completed hard − gene
completed hard − mRNA
completed hard − exon
completed hard − CDS
completed hard − 5' UTR
completed hard − 3' UTR
completed soft − gene
completed soft − mRNA
completed soft − exon
completed soft − CDS
completed soft − 5' UTR
completed soft − 3' UTR
partial hard − gene
partial hard − mRNA
partial hard − exon
partial hard − CDS
partial hard − 5' UTR
partial hard − 3' UTR
partial soft − gene
partial soft − mRNA
partial soft − exon
partial soft − CDS
partial soft − 5' UTR
partial soft − 3' UTR
0
1
2
3
4
enrichment

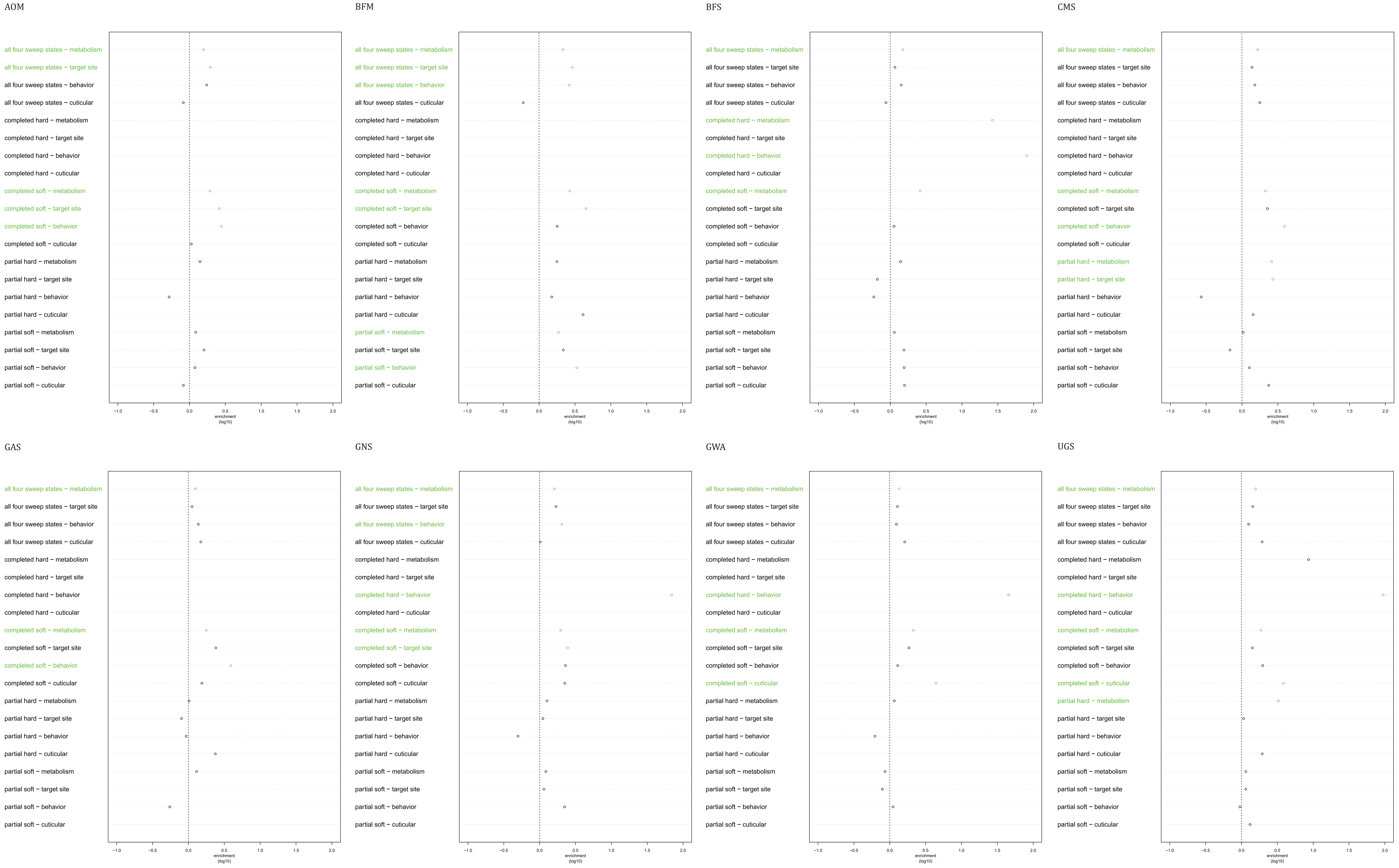
AOM
BFM
BFS
CMS
GAS
GNS
GWA
UGS
all four sweep states − metabolism
all four sweep states − target site
all four sweep states − behavior
all four sweep states − cuticular
completed hard − metabolism
completed hard − target site
completed hard − behavior
completed hard − cuticular
completed soft − metabolism
completed soft − target site
completed soft − behavior
completed soft − cuticular
partial hard − metabolism
partial hard − target site
partial hard − behavior
partial hard − cuticular
partial soft − metabolism
partial soft − target site
partial soft − behavior
partial soft − cuticular
−1.0
−0.5
0.0
0.5
1.0
1.5
2.0
enrichment
(log10)
