## Supplemental Tables (S1-S5) for "Discovery of ongoing selective sweeps within *Anopheles* mosquito populations using deep learning"

**Table S1.** Accuracy rate for deep learning *in silico* test per simulated population dataset.

| **Population** | **Accuracy** |
|---|---|
| AOM | 0.610 |
| BFM | 0.661 |
| BFS | 0.691 |
| CMS | 0.711 |
| GAS | 0.667 |
| GNS | 0.546 |
| GWA | 0.625 |
| UGS | 0.678 |
| *median* | 0.664 |
| *mean* | 0.648 |

**Table S2.** Accuracy rate for five-state deep learning *in silico* test per simulated population dataset (neutral regions and completed sweeps only).

| Population | Accuracy |
|---|---|
| AOM | 0.822 |
| BFM | 0.872 |
| BFS | 0.881 |
| CMS | 0.897 |
| GAS | 0.830 |
| GNS | 0.792 |
| GWA | 0.780 |
| UGS | 0.906 |
| *median* | 0.851 |
| *mean* | 0.848 |

**Table S3.** Nine-state calls for individual chromosome arms and genome-wide for each empirical mosquito population dataset.

| | AOM | BFM | BFS | CMS | GAS | GNS | GWA | UGS | *median* | *mean* |
|---|---|---|---|---|---|---|---|---|---|---|
| | ***neutral*** *(proportion)* | | | | | | | | | |
| **2L** | 0.4524 | 0.6649 | 0.2929 | 0.0167 | 0.3153 | 0.2960 | 0.0395 | 0.1990 | 0.2944 | 0.2846 |
| **2R** | 0.4217 | 0.3822 | 0.2899 | 0.0046 | 0.2227 | 0.3465 | 0.0335 | 0.2231 | 0.2565 | 0.2405 |
| **3L** | 0.5452 | 0.6728 | 0.4012 | 0.0067 | 0.2709 | 0.5540 | 0.3315 | 0.3663 | 0.3838 | 0.3936 |
| **3R** | 0.5181 | 0.6202 | 0.3123 | 0.0069 | 0.2676 | 0.4124 | 0.2433 | 0.3327 | 0.3225 | 0.3392 |
| **genome-wide** | 0.4767 | 0.5608 | 0.3172 | 0.0081 | 0.2626 | 0.3912 | 0.1483 | 0.2761 | 0.2967 | 0.3051 |
| | ***completed hard*** *(proportion)* | | | | | | | | | |
| **2L** | 0.0000 | 0.0000 | 0.0000 | 0.0006 | 0.0004 | 0.0000 | 0.0000 | 0.0004 | 0.0000 | 0.0002 |
| **2R** | 0.0010 | 0.0004 | 0.0002 | 0.0000 | 0.0002 | 0.0002 | 0.0009 | 0.0002 | 0.0002 | 0.0004 |
| **3L** | 0.0000 | 0.0004 | 0.0000 | 0.0000 | 0.0000 | 0.0000 | 0.0008 | 0.0000 | 0.0000 | 0.0002 |
| **3R** | 0.0000 | 0.0008 | 0.0005 | 0.0000 | 0.0005 | 0.0003 | 0.0003 | 0.0007 | 0.0004 | 0.0004 |
| **genome-wide** | 0.0003 | 0.0004 | 0.0002 | 0.0001 | 0.0003 | 0.0002 | 0.0005 | 0.0004 | 0.0003 | 0.0003 |
| | ***completed hard – linked*** *(proportion)* | | | | | | | | | |
| **2L** | 0.0004 | 0.0004 | 0.0017 | 0.0035 | 0.0000 | 0.0019 | 0.0000 | 0.0007 | 0.0006 | 0.0011 |
| **2R** | 0.0045 | 0.0018 | 0.0013 | 0.0034 | 0.0016 | 0.0009 | 0.0018 | 0.0024 | 0.0018 | 0.0022 |
| **3L** | 0.0010 | 0.0020 | 0.0004 | 0.0027 | 0.0005 | 0.0013 | 0.0004 | 0.0008 | 0.0009 | 0.0011 |
| **3R** | 0.0028 | 0.0025 | 0.0020 | 0.0007 | 0.0003 | 0.0000 | 0.0000 | 0.0012 | 0.0009 | 0.0012 |
| **genome-wide** | 0.0026 | 0.0017 | 0.0014 | 0.0025 | 0.0007 | 0.0009 | 0.0007 | 0.0014 | 0.0014 | 0.0015 |
| | ***completed soft*** *(proportion)* | | | | | | | | | |
| **2L** | 0.0587 | 0.0309 | 0.0187 | 0.0273 | 0.0393 | 0.0596 | 0.0665 | 0.0540 | 0.0466 | 0.0444 |
| **2R** | 0.0763 | 0.0307 | 0.0254 | 0.0259 | 0.0664 | 0.0580 | 0.0817 | 0.0647 | 0.0614 | 0.0536 |
| **3L** | 0.0558 | 0.0242 | 0.0234 | 0.0403 | 0.0332 | 0.0511 | 0.0676 | 0.0547 | 0.0457 | 0.0438 |
| **3R** | 0.0446 | 0.0257 | 0.0168 | 0.0329 | 0.0304 | 0.0520 | 0.0716 | 0.0522 | 0.0388 | 0.0408 |
| **genome-wide** | 0.0601 | 0.0281 | 0.0212 | 0.0310 | 0.0449 | 0.0554 | 0.0732 | 0.0571 | 0.0501 | 0.0464 |
| | ***completed soft – linked*** *(proportion)* | | | | | | | | | |
| **2L** | 0.1106 | 0.0872 | 0.1333 | 0.2428 | 0.1537 | 0.3515 | 0.3744 | 0.2050 | 0.1793 | 0.2073 |
| **2R** | 0.1267 | 0.1635 | 0.1310 | 0.2699 | 0.2286 | 0.2983 | 0.4573 | 0.1666 | 0.1976 | 0.2302 |
| **3L** | 0.0837 | 0.0890 | 0.1257 | 0.3544 | 0.1034 | 0.2003 | 0.2353 | 0.1480 | 0.1368 | 0.1675 |
| **3R** | 0.0823 | 0.0920 | 0.1310 | 0.3211 | 0.0783 | 0.2407 | 0.2631 | 0.1572 | 0.1441 | 0.1707 |
| **genome-wide** | 0.1033 | 0.1139 | 0.1305 | 0.2955 | 0.1487 | 0.2755 | 0.3448 | 0.1682 | 0.1584 | 0.1975 |
| | ***partial hard*** *(proportion)* | | | | | | | | | |
| **2L** | 0.0281 | 0.0103 | 0.0463 | 0.0331 | 0.1380 | 0.0331 | 0.0395 | 0.0427 | 0.0363 | 0.0464 |
| **2R** | 0.0363 | 0.0241 | 0.0554 | 0.0836 | 0.0917 | 0.0359 | 0.0276 | 0.0293 | 0.0361 | 0.0480 |
| **3L** | 0.0168 | 0.0071 | 0.0138 | 0.0113 | 0.1624 | 0.0074 | 0.0118 | 0.0119 | 0.0118 | 0.0303 |
| **3R** | 0.0237 | 0.0135 | 0.0388 | 0.0176 | 0.1676 | 0.0315 | 0.0193 | 0.0113 | 0.0215 | 0.0404 |
| **genome-wide** | 0.0277 | 0.0151 | 0.0412 | 0.0405 | 0.1350 | 0.0291 | 0.0248 | 0.0236 | 0.0284 | 0.0422 |
| | ***partial hard – linked*** *(proportion)* | | | | | | | | | |
| **2L** | 0.0768 | 0.0223 | 0.1382 | 0.0864 | 0.1325 | 0.0581 | 0.0512 | 0.0823 | 0.0796 | 0.0810 |

| | | | | | | | | | | |
|---|---|---|---|---|---|---|---|---|---|---|
| **2R** | 0.1011 | 0.0767 | 0.1338 | 0.1535 | 0.1334 | 0.0882 | 0.0372 | 0.0533 | 0.0947 | 0.0971 |
| **3L** | 0.0635 | 0.0115 | 0.0461 | 0.0329 | 0.1867 | 0.0234 | 0.0248 | 0.0184 | 0.0289 | 0.0509 |
| **3R** | 0.0811 | 0.0353 | 0.1112 | 0.0652 | 0.1821 | 0.0746 | 0.0341 | 0.0204 | 0.0699 | 0.0755 |
| **genome-wide** | 0.0841 | 0.0419 | 0.1122 | 0.0914 | 0.1564 | 0.0670 | 0.0370 | 0.0433 | 0.0755 | 0.0791 |
| | ***partial soft*** *(proportion)* | | | | | | | | | |
| **2L** | 0.0885 | 0.0475 | 0.0598 | 0.1040 | 0.0531 | 0.0510 | 0.1170 | 0.1146 | 0.0741 | 0.0794 |
| **2R** | 0.0708 | 0.0504 | 0.0737 | 0.0535 | 0.0499 | 0.0426 | 0.1024 | 0.1292 | 0.0621 | 0.0716 |
| **3L** | 0.0659 | 0.0461 | 0.0734 | 0.0839 | 0.0538 | 0.0438 | 0.1105 | 0.1296 | 0.0696 | 0.0759 |
| **3R** | 0.0761 | 0.0473 | 0.0702 | 0.1547 | 0.0545 | 0.0426 | 0.1249 | 0.1447 | 0.0732 | 0.0894 |
| **genome-wide** | 0.0751 | 0.0481 | 0.0698 | 0.0986 | 0.0526 | 0.0445 | 0.1133 | 0.1308 | 0.0724 | 0.0791 |
| | ***partial soft – linked*** *(proportion)* | | | | | | | | | |
| **2L** | 0.1846 | 0.1365 | 0.3092 | 0.4855 | 0.1678 | 0.1489 | 0.3119 | 0.3013 | 0.2429 | 0.2557 |
| **2R** | 0.1616 | 0.2704 | 0.2893 | 0.4055 | 0.2054 | 0.1293 | 0.2575 | 0.3312 | 0.2640 | 0.2563 |
| **3L** | 0.1683 | 0.1469 | 0.3159 | 0.4679 | 0.1891 | 0.1188 | 0.2172 | 0.2704 | 0.2031 | 0.2368 |
| **3R** | 0.1712 | 0.1629 | 0.3173 | 0.4010 | 0.2186 | 0.1459 | 0.2436 | 0.2795 | 0.2311 | 0.2425 |
| **genome-wide** | 0.1702 | 0.1898 | 0.3062 | 0.4322 | 0.1989 | 0.1362 | 0.2575 | 0.2991 | 0.2282 | 0.2487 |
| | ***neutral*** *(count)* | | | | | | | | | |
| **2L** | 1,125 | 1,875 | 848 | 52 | 802 | 795 | 108 | 568 | 798.5 | 771.6 |
| **2R** | 1,764 | 1,730 | 1,361 | 23 | 950 | 1,552 | 147 | 1,034 | 1,197.5 | 1,070.1 |
| **3L** | 1,134 | 1,694 | 1,044 | 20 | 579 | 1,278 | 789 | 958 | 1,001.0 | 937.0 |
| **3R** | 1,858 | 2,441 | 1,264 | 31 | 977 | 1,569 | 935 | 1,352 | 1,308.0 | 1,303.4 |
| **genome-wide** | 5,881 | 7,740 | 4,517 | 126 | 3,308 | 5,194 | 1,979 | 3,912 | 4,214.5 | 4,082.1 |
| | ***completed hard*** *(count)* | | | | | | | | | |
| **2L** | 0 | 0 | 0 | 2 | 1 | 0 | 0 | 1 | 0.0 | 0.5 |
| **2R** | 4 | 2 | 1 | 0 | 1 | 1 | 4 | 1 | 1.0 | 1.8 |
| **3L** | 0 | 1 | 0 | 0 | 0 | 0 | 2 | 0 | 0.0 | 0.4 |
| **3R** | 0 | 3 | 2 | 0 | 2 | 1 | 1 | 3 | 1.5 | 1.5 |
| **genome-wide** | 4 | 6 | 3 | 2 | 4 | 2 | 7 | 5 | 4.0 | 4.1 |
| | ***completed hard – linked*** *(count)* | | | | | | | | | |
| **2L** | 1 | 1 | 5 | 11 | 0 | 5 | 0 | 2 | 1.5 | 3.1 |
| **2R** | 19 | 8 | 6 | 17 | 7 | 4 | 8 | 11 | 8.0 | 10.0 |
| **3L** | 2 | 5 | 1 | 8 | 1 | 3 | 1 | 2 | 2.0 | 2.9 |
| **3R** | 10 | 10 | 8 | 3 | 1 | 0 | 0 | 5 | 4.0 | 4.6 |
| **genome-wide** | 32 | 24 | 20 | 39 | 9 | 12 | 9 | 20 | 20.0 | 20.6 |
| | ***completed soft*** *(count)* | | | | | | | | | |
| **2L** | 146 | 87 | 54 | 85 | 100 | 160 | 182 | 154 | 123.0 | 121.0 |
| **2R** | 319 | 139 | 119 | 129 | 283 | 260 | 358 | 300 | 271.5 | 238.4 |
| **3L** | 116 | 61 | 61 | 121 | 71 | 118 | 161 | 143 | 117.0 | 106.5 |
| **3R** | 160 | 101 | 68 | 148 | 111 | 198 | 275 | 212 | 154.0 | 159.1 |
| **genome-wide** | 741 | 388 | 302 | 483 | 565 | 736 | 976 | 809 | 650.5 | 625.0 |
| | ***completed soft – linked*** *(count)* | | | | | | | | | |
| **2L** | 275 | 246 | 386 | 756 | 391 | 944 | 1,024 | 585 | 488.0 | 575.9 |

| | | | | | | | | | | |
|---|---|---|---|---|---|---|---|---|---|---|
| **2R** | 530 | 740 | 615 | 1,343 | 975 | 1,336 | 2,005 | 772 | 873.5 | 1,039.5 |
| **3L** | 174 | 224 | 327 | 1,065 | 221 | 462 | 560 | 387 | 357.0 | 427.5 |
| **3R** | 295 | 362 | 530 | 1,443 | 286 | 916 | 1,011 | 639 | 584.5 | 685.3 |
| **genome-wide** | 1,274 | 1,572 | 1,858 | 4,607 | 1,873 | 3,658 | 4,600 | 2,383 | 2,128.0 | 2,728.1 |
| | ***partial hard*** *(count)* | | | | | | | | | |
| **2L** | 70 | 29 | 134 | 103 | 351 | 89 | 108 | 122 | 105.5 | 125.8 |
| **2R** | 152 | 109 | 260 | 416 | 391 | 161 | 121 | 136 | 156.5 | 218.3 |
| **3L** | 35 | 18 | 36 | 34 | 347 | 17 | 28 | 31 | 32.5 | 68.3 |
| **3R** | 85 | 53 | 157 | 79 | 612 | 120 | 74 | 46 | 82.0 | 153.3 |
| **genome-wide** | 342 | 209 | 587 | 632 | 1,701 | 387 | 331 | 335 | 364.5 | 565.5 |
| | ***partial hard – linked*** *(count)* | | | | | | | | | |
| **2L** | 191 | 63 | 400 | 269 | 337 | 156 | 140 | 235 | 213.0 | 223.9 |
| **2R** | 423 | 347 | 628 | 764 | 569 | 395 | 163 | 247 | 409.0 | 442.0 |
| **3L** | 132 | 29 | 120 | 99 | 399 | 54 | 59 | 48 | 79.0 | 117.5 |
| **3R** | 291 | 139 | 450 | 293 | 665 | 284 | 131 | 83 | 287.5 | 292.0 |
| **genome-wide** | 1,037 | 578 | 1,598 | 1,425 | 1,970 | 889 | 493 | 613 | 963.0 | 1,075.4 |
| | ***partial soft*** *(count)* | | | | | | | | | |
| **2L** | 220 | 134 | 173 | 324 | 135 | 137 | 320 | 327 | 196.5 | 221.3 |
| **2R** | 296 | 228 | 346 | 266 | 213 | 191 | 449 | 599 | 281.0 | 323.5 |
| **3L** | 137 | 116 | 191 | 252 | 115 | 101 | 263 | 339 | 164.0 | 189.3 |
| **3R** | 273 | 186 | 284 | 695 | 199 | 162 | 480 | 588 | 278.5 | 358.4 |
| **genome-wide** | 926 | 664 | 994 | 1,537 | 662 | 591 | 1,512 | 1,853 | 960.0 | 1,092.4 |
| | ***partial soft – linked*** *(count)* | | | | | | | | | |
| **2L** | 459 | 385 | 895 | 1,512 | 427 | 400 | 853 | 860 | 656.0 | 723.9 |
| **2R** | 676 | 1,224 | 1,358 | 2,018 | 876 | 579 | 1,129 | 1,535 | 1,176.5 | 1,174.4 |
| **3L** | 350 | 370 | 822 | 1,406 | 404 | 274 | 517 | 707 | 460.5 | 606.3 |
| **3R** | 614 | 641 | 1,284 | 1,802 | 798 | 555 | 936 | 1,136 | 867.0 | 970.8 |
| **genome-wide** | 2,099 | 2,620 | 4,359 | 6,738 | 2,505 | 1,808 | 3,435 | 4,238 | 3,027.5 | 3,475.3 |
| | ***neutral*** *(proportion)* *corrected for false discovery* | | | | | | | | | |
| **2L** | 0.4922 | 0.7173 | 0.3247 | 0.0387 | 0.3700 | 0.3486 | 0.0502 | 0.2369 | 0.3367 | 0.3223 |
| **2R** | 0.4589 | 0.4122 | 0.3214 | 0.0107 | 0.2614 | 0.4081 | 0.0427 | 0.2656 | 0.2935 | 0.2726 |
| **3L** | 0.5932 | 0.7257 | 0.4448 | 0.0154 | 0.3180 | 0.6525 | 0.4218 | 0.4361 | 0.4405 | 0.4510 |
| **3R** | 0.5638 | 0.6690 | 0.3463 | 0.0160 | 0.3141 | 0.4857 | 0.3095 | 0.3960 | 0.3712 | 0.3876 |
| **genome-wide** | 0.5188 | 0.6050 | 0.3517 | 0.0188 | 0.3082 | 0.4608 | 0.1887 | 0.3287 | 0.3402 | 0.3476 |
| | ***completed hard*** *(proportion)* *corrected for false discovery* | | | | | | | | | |
| **2L** | 0.0000 | 0.0000 | 0.0000 | 0.0006 | 0.0004 | 0.0000 | 0.0000 | 0.0004 | 0.0000 | 0.0002 |
| **2R** | 0.0010 | 0.0004 | 0.0002 | 0.0000 | 0.0002 | 0.0002 | 0.0009 | 0.0002 | 0.0002 | 0.0004 |
| **3L** | 0.0000 | 0.0004 | 0.0000 | 0.0000 | 0.0000 | 0.0000 | 0.0008 | 0.0000 | 0.0000 | 0.0002 |
| **3R** | 0.0000 | 0.0008 | 0.0005 | 0.0000 | 0.0005 | 0.0003 | 0.0003 | 0.0007 | 0.0004 | 0.0004 |
| **genome-wide** | 0.0003 | 0.0004 | 0.0002 | 0.0001 | 0.0003 | 0.0002 | 0.0005 | 0.0004 | 0.0003 | 0.0003 |

| | | | | | | | | | | |
|---|---|---|---|---|---|---|---|---|---|---|
| | ***completed hard – linked*** *(proportion)* *corrected for false discovery* | | | | | | | | | |
| **2L** | 0.0004 | -0.0011 | 0.0017 | 0.0035 | 0.0000 | 0.0019 | 0.0000 | 0.0007 | 0.0006 | 0.0009 |
| **2R** | 0.0045 | 0.0009 | 0.0013 | 0.0034 | 0.0016 | 0.0009 | 0.0018 | 0.0024 | 0.0017 | 0.0021 |
| **3L** | 0.0010 | 0.0005 | 0.0004 | 0.0027 | 0.0005 | 0.0013 | 0.0004 | 0.0008 | 0.0006 | 0.0009 |
| **3R** | 0.0028 | 0.0012 | 0.0020 | 0.0007 | 0.0003 | 0.0000 | 0.0000 | 0.0012 | 0.0009 | 0.0010 |
| **genome-wide** | 0.0026 | 0.0005 | 0.0014 | 0.0025 | 0.0007 | 0.0009 | 0.0007 | 0.0014 | 0.0012 | 0.0013 |
| | ***completed soft*** *(proportion)* *corrected for false discovery* | | | | | | | | | |
| **2L** | 0.0498 | 0.0258 | 0.0177 | 0.0267 | 0.0345 | 0.0495 | 0.0652 | 0.0485 | 0.0415 | 0.0397 |
| **2R** | 0.0680 | 0.0278 | 0.0244 | 0.0258 | 0.0630 | 0.0462 | 0.0805 | 0.0586 | 0.0524 | 0.0493 |
| **3L** | 0.0451 | 0.0191 | 0.0221 | 0.0400 | 0.0291 | 0.0322 | 0.0563 | 0.0447 | 0.0361 | 0.0361 |
| **3R** | 0.0345 | 0.0210 | 0.0158 | 0.0327 | 0.0263 | 0.0380 | 0.0632 | 0.0431 | 0.0336 | 0.0343 |
| **genome-wide** | 0.0507 | 0.0239 | 0.0202 | 0.0307 | 0.0408 | 0.0421 | 0.0681 | 0.0495 | 0.0415 | 0.0407 |
| | ***completed soft – linked*** *(proportion)* *corrected for false discovery* | | | | | | | | | |
| **2L** | 0.1012 | 0.0822 | 0.1304 | 0.2419 | 0.1463 | 0.3396 | 0.3727 | 0.2036 | 0.1749 | 0.2022 |
| **2R** | 0.1180 | 0.1606 | 0.1281 | 0.2697 | 0.2234 | 0.2844 | 0.4559 | 0.1650 | 0.1942 | 0.2256 |
| **3L** | 0.0724 | 0.0839 | 0.1217 | 0.3541 | 0.0971 | 0.1781 | 0.2214 | 0.1454 | 0.1335 | 0.1592 |
| **3R** | 0.0716 | 0.0873 | 0.1278 | 0.3207 | 0.0721 | 0.2242 | 0.2529 | 0.1549 | 0.1414 | 0.1639 |
| **genome-wide** | 0.0934 | 0.1097 | 0.1273 | 0.2951 | 0.1425 | 0.2598 | 0.3385 | 0.1662 | 0.1544 | 0.1916 |
| | ***partial hard*** *(proportion)* *corrected for false discovery* | | | | | | | | | |
| **2L** | 0.0277 | 0.0103 | 0.0460 | 0.0330 | 0.1369 | 0.0331 | 0.0395 | 0.0425 | 0.0363 | 0.0461 |
| **2R** | 0.0359 | 0.0241 | 0.0551 | 0.0836 | 0.0909 | 0.0359 | 0.0276 | 0.0291 | 0.0359 | 0.0478 |
| **3L** | 0.0162 | 0.0071 | 0.0134 | 0.0113 | 0.1614 | 0.0074 | 0.0118 | 0.0114 | 0.0116 | 0.0300 |
| **3R** | 0.0231 | 0.0135 | 0.0384 | 0.0175 | 0.1667 | 0.0315 | 0.0193 | 0.0109 | 0.0212 | 0.0401 |
| **genome-wide** | 0.0272 | 0.0151 | 0.0409 | 0.0405 | 0.1341 | 0.0291 | 0.0248 | 0.0233 | 0.0282 | 0.0419 |
| | ***partial hard – linked*** *(proportion)* *corrected for false discovery* | | | | | | | | | |
| **2L** | 0.0748 | 0.0223 | 0.1378 | 0.0863 | 0.1325 | 0.0567 | 0.0512 | 0.0821 | 0.0785 | 0.0805 |
| **2R** | 0.0993 | 0.0767 | 0.1335 | 0.1535 | 0.1334 | 0.0866 | 0.0372 | 0.0530 | 0.0929 | 0.0966 |
| **3L** | 0.0611 | 0.0115 | 0.0457 | 0.0329 | 0.1867 | 0.0208 | 0.0248 | 0.0179 | 0.0289 | 0.0502 |
| **3R** | 0.0789 | 0.0353 | 0.1108 | 0.0652 | 0.1821 | 0.0727 | 0.0341 | 0.0200 | 0.0689 | 0.0749 |
| **genome-wide** | 0.0820 | 0.0419 | 0.1119 | 0.0914 | 0.1564 | 0.0651 | 0.0370 | 0.0429 | 0.0736 | 0.0786 |
| | ***partial soft*** *(proportion)* *corrected for false discovery* | | | | | | | | | |
| **2L** | 0.0816 | 0.0274 | 0.0533 | 0.1026 | 0.0468 | 0.0385 | 0.1127 | 0.0987 | 0.0674 | 0.0702 |
| **2R** | 0.0643 | 0.0388 | 0.0673 | 0.0530 | 0.0455 | 0.0280 | 0.0987 | 0.1114 | 0.0587 | 0.0634 |
| **3L** | 0.0576 | 0.0257 | 0.0645 | 0.0833 | 0.0484 | 0.0203 | 0.0742 | 0.1004 | 0.0610 | 0.0593 |
| **3R** | 0.0682 | 0.0285 | 0.0632 | 0.1540 | 0.0492 | 0.0251 | 0.0983 | 0.1182 | 0.0657 | 0.0756 |
| **genome-wide** | 0.0678 | 0.0312 | 0.0628 | 0.0979 | 0.0473 | 0.0279 | 0.0971 | 0.1088 | 0.0653 | 0.0676 |
| | ***partial soft – linked*** *(proportion)* *corrected for false discovery* | | | | | | | | | |
| **2L** | 0.1723 | 0.1157 | 0.2884 | 0.4666 | 0.1327 | 0.1322 | 0.3085 | 0.2866 | 0.2294 | 0.2379 |

| | | | | | | | | | | |
|---|---|---|---|---|---|---|---|---|---|---|
| **2R** | 0.1501 | 0.2584 | 0.2687 | 0.4003 | 0.1806 | 0.1097 | 0.2546 | 0.3147 | 0.2565 | 0.2421 |
| **3L** | 0.1534 | 0.1259 | 0.2874 | 0.4603 | 0.1588 | 0.0874 | 0.1885 | 0.2433 | 0.1737 | 0.2132 |
| **3R** | 0.1571 | 0.1435 | 0.2951 | 0.3931 | 0.1887 | 0.1225 | 0.2225 | 0.2550 | 0.2056 | 0.2222 |
| **genome-wide** | 0.1572 | 0.1723 | 0.2836 | 0.4230 | 0.1696 | 0.1141 | 0.2446 | 0.2787 | 0.2085 | 0.2304 |
| | ***neutral*** *(count)* *corrected for false discovery* | | | | | | | | | |
| **2L** | 1,224.2 | 2,022.7 | 940.1 | 120.6 | 941.3 | 936.4 | 137.4 | 676.2 | 938.3 | 874.9 |
| **2R** | 1,919.5 | 1,866.2 | 1,508.9 | 53.4 | 1,115.0 | 1,828.0 | 187.0 | 1,231.0 | 1,369.9 | 1,213.6 |
| **3L** | 1,233.9 | 1,827.4 | 1,157.4 | 46.4 | 679.6 | 1,505.3 | 1,003.8 | 1,140.5 | 1,149.0 | 1,074.3 |
| **3R** | 2,021.8 | 2,633.2 | 1,401.3 | 71.9 | 1,146.7 | 1,848.1 | 1,189.6 | 1,609.5 | 1,505.4 | 1,490.3 |
| **genome-wide** | 6,399.3 | 8,349.5 | 5,007.8 | 292.3 | 3,882.6 | 6,117.8 | 2,517.8 | 4,657.1 | 4,832.5 | 4,653.0 |
| | ***completed hard*** *(count)* *corrected for false discovery* | | | | | | | | | |
| **2L** | 0.0 | 0.0 | 0.0 | 1.9 | 1.0 | 0.0 | 0.0 | 1.0 | 0.0 | 0.5 |
| **2R** | 4.0 | 2.0 | 1.0 | -0.1 | 1.0 | 1.0 | 4.0 | 1.0 | 1.0 | 1.7 |
| **3L** | 0.0 | 1.0 | 0.0 | 0.0 | 0.0 | 0.0 | 2.0 | 0.0 | 0.0 | 0.4 |
| **3R** | 0.0 | 3.0 | 2.0 | -0.1 | 2.0 | 1.0 | 1.0 | 3.0 | 1.5 | 1.5 |
| **genome-wide** | 4.0 | 6.0 | 3.0 | 1.7 | 4.0 | 2.0 | 7.0 | 5.0 | 4.0 | 4.1 |
| | ***completed hard – linked*** *(count)* *corrected for false discovery* | | | | | | | | | |
| **2L** | 1.0 | -3.0 | 5.0 | 11.0 | 0.0 | 5.0 | 0.0 | 2.0 | 1.5 | 2.6 |
| **2R** | 19.0 | 4.3 | 6.0 | 17.0 | 7.0 | 4.0 | 8.0 | 11.0 | 7.5 | 9.5 |
| **3L** | 2.0 | 1.3 | 1.0 | 8.0 | 1.0 | 3.0 | 1.0 | 2.0 | 1.7 | 2.4 |
| **3R** | 10.0 | 4.7 | 8.0 | 3.0 | 1.0 | 0.0 | 0.0 | 5.0 | 3.9 | 4.0 |
| **genome-wide** | 32.0 | 7.3 | 20.0 | 39.0 | 9.0 | 12.0 | 9.0 | 20.0 | 16.0 | 18.5 |
| | ***completed soft*** *(count)* *corrected for false discovery* | | | | | | | | | |
| **2L** | 124.0 | 72.8 | 51.2 | 83.2 | 87.8 | 132.8 | 178.3 | 138.4 | 105.9 | 108.6 |
| **2R** | 284.4 | 125.9 | 114.5 | 128.2 | 268.5 | 207.0 | 353.0 | 271.7 | 237.7 | 219.1 |
| **3L** | 93.8 | 48.2 | 57.5 | 120.3 | 62.2 | 74.3 | 133.9 | 116.8 | 84.1 | 88.4 |
| **3R** | 123.6 | 82.6 | 63.8 | 146.9 | 96.1 | 144.4 | 242.9 | 175.0 | 134.0 | 134.4 |
| **genome-wide** | 625.8 | 329.6 | 287.0 | 478.6 | 514.5 | 558.6 | 908.0 | 701.9 | 536.6 | 550.5 |
| | ***completed soft – linked*** *(count)* *corrected for false discovery* | | | | | | | | | |
| **2L** | 251.7 | 231.8 | 377.5 | 753.3 | 372.2 | 912.2 | 1,019.5 | 580.9 | 479.2 | 562.4 |
| **2R** | 493.5 | 726.9 | 601.4 | 1,341.8 | 952.7 | 1,273.8 | 1,998.8 | 764.6 | 858.7 | 1,019.2 |
| **3L** | 150.6 | 211.2 | 316.6 | 1,064.0 | 207.4 | 410.8 | 526.9 | 380.2 | 348.4 | 408.4 |
| **3R** | 256.6 | 343.6 | 517.4 | 1,441.4 | 263.1 | 853.2 | 971.7 | 629.3 | 573.4 | 659.5 |
| **genome-wide** | 1,152.4 | 1,513.6 | 1,812.9 | 4,600.6 | 1,795.3 | 3,450.0 | 4,516.9 | 2,355.1 | 2,084.0 | 2,649.6 |
| | ***partial hard*** *(count)* *corrected for false discovery* | | | | | | | | | |
| **2L** | 68.8 | 29.0 | 133.1 | 102.8 | 348.2 | 89.0 | 108.0 | 121.3 | 105.4 | 125.0 |
| **2R** | 150.1 | 109.0 | 258.5 | 415.9 | 387.7 | 161.0 | 121.0 | 134.8 | 155.5 | 217.2 |
| **3L** | 33.8 | 18.0 | 34.8 | 33.9 | 345.0 | 17.0 | 28.0 | 29.9 | 31.8 | 67.5 |

| | | | | | | | | | | |
|---|---|---|---|---|---|---|---|---|---|---|
| **3R** | 83.0 | 53.0 | 155.6 | 78.9 | 608.6 | 120.0 | 74.0 | 44.4 | 80.9 | 152.2 |
| **genome-wide** | 335.6 | 209.0 | 582.0 | 631.4 | 1,689.4 | 387.0 | 331.0 | 330.3 | 361.3 | 562.0 |
| | ***partial hard – linked*** *(count)* *corrected for false discovery* | | | | | | | | | |
| **2L** | 186.1 | 63.0 | 399.1 | 268.9 | 337.0 | 152.3 | 140.0 | 234.3 | 210.2 | 222.6 |
| **2R** | 415.3 | 347.0 | 626.5 | 763.9 | 569.0 | 387.7 | 163.0 | 245.8 | 401.5 | 439.8 |
| **3L** | 127.1 | 29.0 | 118.8 | 99.0 | 399.0 | 48.0 | 59.0 | 46.9 | 79.0 | 115.8 |
| **3R** | 282.9 | 139.0 | 448.6 | 292.9 | 665.0 | 276.6 | 131.0 | 81.4 | 279.8 | 289.7 |
| **genome-wide** | 1,011.4 | 578.0 | 1,593.0 | 1,424.7 | 1,970.0 | 864.5 | 493.0 | 608.3 | 938.0 | 1,067.9 |
| | ***partial soft*** *(count)* *corrected for false discovery* | | | | | | | | | |
| **2L** | 202.9 | 77.4 | 154.2 | 319.4 | 119.0 | 103.3 | 308.2 | 281.7 | 178.5 | 195.8 |
| **2R** | 269.1 | 175.7 | 315.8 | 264.0 | 194.0 | 125.2 | 432.9 | 516.5 | 266.5 | 286.7 |
| **3L** | 119.7 | 64.8 | 167.9 | 250.2 | 103.4 | 46.8 | 176.7 | 262.6 | 143.8 | 149.0 |
| **3R** | 244.7 | 112.3 | 256.0 | 692.3 | 179.5 | 95.5 | 377.7 | 480.2 | 250.3 | 304.8 |
| **genome-wide** | 836.4 | 430.2 | 893.8 | 1,525.9 | 596.0 | 370.8 | 1,295.5 | 1,541.0 | 865.1 | 936.2 |
| | ***partial soft – linked*** *(count)* *corrected for false discovery* | | | | | | | | | |
| **2L** | 428.4 | 326.3 | 834.8 | 1,452.9 | 337.6 | 355.1 | 843.7 | 818.1 | 623.2 | 674.6 |
| **2R** | 628.0 | 1,169.9 | 1,261.4 | 1,991.9 | 770.1 | 491.3 | 1,116.3 | 1,458.7 | 1,143.1 | 1,110.9 |
| **3L** | 319.2 | 317.0 | 747.9 | 1,383.3 | 339.4 | 201.7 | 448.7 | 636.3 | 394.1 | 549.2 |
| **3R** | 563.5 | 564.6 | 1,194.3 | 1,766.8 | 689.1 | 466.3 | 855.1 | 1,036.2 | 772.1 | 892.0 |
| **genome-wide** | 1,939.0 | 2,377.9 | 4,038.5 | 6,594.8 | 2,136.2 | 1,514.3 | 3,263.8 | 3,949.3 | 2,820.8 | 3,226.7 |

**Table S4.** Enrichments of selective sweeps within DNA regions for each empirical mosquito population dataset.

| | | AOM | BFM | BFS | CMS | GAS | GNS | GWA | UGS |
|---|---|---|---|---|---|---|---|---|---|
| | | ***all four sweep states*** | | | | | | | |
| **gene** | **count** | 1177 | 820 | 1173 | 1530 | 1636 | 1028 | 1698 | 1839 |
| | **enrichment** | 1.11 | 1.24 | 1.19 | 1.13 | 1.08 | 1.14 | 1.15 | 1.18 |
| | ***p*-value** | ≤0.0001 | ≤0.0001 | ≤0.0001 | ≤0.0001 | ≤0.0001 | ≤0.0001 | ≤0.0001 | ≤0.0001 |
| **mRNA** | **count** | 1176 | 819 | 1171 | 1529 | 1634 | 1028 | 1697 | 1836 |
| | **enrichment** | 1.11 | 1.24 | 1.19 | 1.13 | 1.08 | 1.14 | 1.15 | 1.18 |
| | ***p*-value** | ≤0.0001 | ≤0.0001 | ≤0.0001 | ≤0.0001 | ≤0.0001 | ≤0.0001 | ≤0.0001 | ≤0.0001 |
| **exon** | **count** | 919 | 700 | 951 | 1230 | 1280 | 824 | 1365 | 1506 |
| | **enrichment** | 1.18 | 1.44 | 1.31 | 1.23 | 1.14 | 1.24 | 1.25 | 1.31 |
| | ***p*-value** | ≤0.0001 | ≤0.0001 | ≤0.0001 | ≤0.0001 | ≤0.0001 | ≤0.0001 | ≤0.0001 | ≤0.0001 |
| **CDS** | **count** | 887 | 685 | 930 | 1198 | 1231 | 798 | 1325 | 1468 |
| | **enrichment** | 1.19 | 1.48 | 1.36 | 1.26 | 1.16 | 1.26 | 1.28 | 1.35 |
| | ***p*-value** | ≤0.0001 | ≤0.0001 | ≤0.0001 | ≤0.0001 | ≤0.0001 | ≤0.0001 | ≤0.0001 | ≤0.0001 |
| **5’ UTR** | **count** | 338 | 265 | 358 | 445 | 446 | 302 | 507 | 537 |
| | **enrichment** | 1.24 | 1.56 | 1.41 | 1.28 | 1.16 | 1.30 | 1.34 | 1.34 |
| | ***p*-value** | ≤0.0001 | ≤0.0001 | ≤0.0001 | ≤0.0001 | ≤0.0001 | ≤0.0001 | ≤0.0001 | ≤0.0001 |
| **3’ UTR** | **count** | 312 | 247 | 343 | 414 | 416 | 278 | 440 | 502 |
| | **enrichment** | 1.34 | 1.70 | 1.59 | 1.38 | 1.26 | 1.40 | 1.37 | 1.47 |
| | ***p*-value** | ≤0.0001 | ≤0.0001 | ≤0.0001 | ≤0.0001 | ≤0.0001 | ≤0.0001 | ≤0.0001 | ≤0.0001 |
| | | ***completed hard*** | | | | | | | |
| **gene** | **count** | 1 | 5 | 2 | 2 | 2 | 1 | 4 | 4 |
| | **enrichment** | 0.45 | 1.68 | 1.30 | 1.71 | 0.93 | 0.95 | 1.15 | 1.51 |
| | ***p*-value** | 0.9278 | 0.1034 | 0.5170 | 0.5043 | 0.7459 | 0.7780 | 0.5054 | 0.2250 |
| **mRNA** | **count** | 1 | 5 | 2 | 2 | 2 | 1 | 4 | 4 |
| | **enrichment** | 0.45 | 1.69 | 1.30 | 1.71 | 0.93 | 0.95 | 1.15 | 1.52 |
| | ***p*-value** | 0.9276 | 0.1020 | 0.5143 | 0.5043 | 0.7436 | 0.7770 | 0.5038 | 0.2237 |
| **exon** | **count** | 1 | 3 | 2 | 2 | 2 | 1 | 4 | 4 |
| | **enrichment** | 0.59 | 1.35 | 1.78 | 2.45 | 1.28 | 1.28 | 1.50 | 2.09 |
| | ***p*-value** | 0.8484 | 0.3910 | 0.3126 | 0.2786 | 0.5070 | 0.6298 | 0.2734 | 0.0762 |
| **CDS** | **count** | 1 | 3 | 2 | 2 | 2 | 1 | 4 | 4 |
| | **enrichment** | 0.62 | 1.42 | 1.87 | 2.61 | 1.35 | 1.34 | 1.58 | 2.21 |
| | ***p*-value** | 0.8288 | 0.3574 | 0.2881 | 0.2549 | 0.4743 | 0.6087 | 0.2387 | 0.0647 |
| **5’ UTR** | **count** | 0 | 2 | 1 | 0 | 1 | 0 | 3 | 1 |
| | **enrichment** | 0.00 | 2.56 | 2.55 | 0.00 | 1.80 | 0.00 | 3.17 | 1.48 |
| | ***p*-value** | 1.0000 | 0.1804 | 0.3424 | 1.0000 | 0.4523 | 1.0000 | 0.0652 | 0.5172 |
| **3’ UTR** | **count** | 0 | 2 | 1 | 0 | 1 | 1 | 0 | 2 |
| | **enrichment** | 0.00 | 2.94 | 2.92 | 0.00 | 2.13 | 4.15 | 0.00 | 3.50 |
| | ***p*-value** | 1.0000 | 0.1430 | 0.3015 | 1.0000 | 0.3906 | 0.2259 | 1.0000 | 0.1017 |
| | | ***completed soft*** | | | | | | | |
| **gene** | **count** | 485 | 345 | 283 | 376 | 425 | 521 | 715 | 617 |
| | **enrichment** | 1.25 | 1.70 | 1.83 | 1.57 | 1.41 | 1.36 | 1.41 | 1.47 |

| | | | | | | | | | |
|---|---|---|---|---|---|---|---|---|---|
| | ***p*-value** | ≤0.0001 | ≤0.0001 | ≤0.0001 | ≤0.0001 | ≤0.0001 | ≤0.0001 | ≤0.0001 | ≤0.0001 |
| **mRNA** | **count** | 485 | 345 | 283 | 376 | 425 | 521 | 715 | 617 |
| | **enrichment** | 1.25 | 1.70 | 1.83 | 1.58 | 1.41 | 1.36 | 1.41 | 1.48 |
| | ***p*-value** | ≤0.0001 | ≤0.0001 | ≤0.0001 | ≤0.0001 | ≤0.0001 | ≤0.0001 | ≤0.0001 | ≤0.0001 |
| **exon** | **count** | 424 | 335 | 276 | 356 | 394 | 459 | 665 | 582 |
| | **enrichment** | 1.47 | 2.24 | 2.39 | 1.99 | 1.76 | 1.62 | 1.77 | 1.88 |
| | ***p*-value** | ≤0.0001 | ≤0.0001 | ≤0.0001 | ≤0.0001 | ≤0.0001 | ≤0.0001 | ≤0.0001 | ≤0.0001 |
| **CDS** | **count** | 416 | 334 | 274 | 351 | 389 | 453 | 653 | 575 |
| | **enrichment** | 1.52 | 2.35 | 2.51 | 2.08 | 1.83 | 1.68 | 1.83 | 1.96 |
| | ***p*-value** | ≤0.0001 | ≤0.0001 | ≤0.0001 | ≤0.0001 | ≤0.0001 | ≤0.0001 | ≤0.0001 | ≤0.0001 |
| **5' UTR** | **count** | 157 | 126 | 96 | 124 | 141 | 172 | 239 | 213 |
| | **enrichment** | 1.55 | 2.40 | 2.37 | 2.01 | 1.78 | 1.74 | 1.83 | 1.96 |
| | ***p*-value** | ≤0.0001 | ≤0.0001 | ≤0.0001 | ≤0.0001 | ≤0.0001 | ≤0.0001 | ≤0.0001 | ≤0.0001 |
| **3' UTR** | **count** | 153 | 131 | 98 | 131 | 137 | 163 | 245 | 208 |
| | **enrichment** | 1.77 | 2.93 | 2.86 | 2.46 | 2.03 | 1.93 | 2.20 | 2.26 |
| | ***p*-value** | ≤0.0001 | ≤0.0001 | ≤0.0001 | ≤0.0001 | ≤0.0001 | ≤0.0001 | ≤0.0001 | ≤0.0001 |
| | | ***partial hard*** | | | | | | | |
| **gene** | **count** | 180 | 99 | 358 | 375 | 845 | 210 | 189 | 206 |
| | **enrichment** | 0.99 | 0.89 | 1.13 | 1.09 | 0.98 | 1.01 | 1.05 | 1.12 |
| | ***p*-value** | 0.5989 | 0.9251 | 0.0018 | 0.0364 | 0.7931 | 0.4632 | 0.2261 | 0.0305 |
| **mRNA** | **count** | 180 | 99 | 357 | 374 | 844 | 210 | 189 | 206 |
| | **enrichment** | 0.99 | 0.89 | 1.13 | 1.08 | 0.98 | 1.01 | 1.05 | 1.12 |
| | ***p*-value** | 0.5896 | 0.9219 | 0.0020 | 0.0392 | 0.7849 | 0.4540 | 0.2215 | 0.0294 |
| **exon** | **count** | 116 | 74 | 250 | 290 | 612 | 141 | 127 | 153 |
| | **enrichment** | 0.86 | 0.90 | 1.08 | 1.13 | 0.96 | 0.92 | 0.96 | 1.13 |
| | ***p*-value** | 0.9707 | 0.8584 | 0.1037 | 0.0141 | 0.8605 | 0.8407 | 0.6916 | 0.0487 |
| **CDS** | **count** | 110 | 69 | 244 | 282 | 584 | 130 | 122 | 150 |
| | **enrichment** | 0.86 | 0.88 | 1.11 | 1.16 | 0.97 | 0.90 | 0.98 | 1.18 |
| | ***p*-value** | 0.9685 | 0.8856 | 0.0415 | 0.0060 | 0.8130 | 0.9043 | 0.6225 | 0.0169 |
| **5' UTR** | **count** | 39 | 30 | 108 | 103 | 210 | 50 | 48 | 53 |
| | **enrichment** | 0.82 | 1.01 | 1.30 | 1.09 | 0.97 | 0.92 | 1.02 | 1.10 |
| | ***p*-value** | 0.9118 | 0.4890 | 0.0046 | 0.2032 | 0.6747 | 0.7255 | 0.4641 | 0.2741 |
| **3' UTR** | **count** | 42 | 26 | 101 | 105 | 189 | 42 | 34 | 50 |
| | **enrichment** | 1.04 | 1.03 | 1.44 | 1.33 | 1.01 | 0.91 | 0.86 | 1.25 |
| | ***p*-value** | 0.4317 | 0.4620 | 0.0003 | 0.0053 | 0.4343 | 0.7325 | 0.8156 | 0.0725 |
| | | ***partial soft*** | | | | | | | |
| **gene** | **count** | 511 | 371 | 530 | 777 | 364 | 296 | 790 | 1012 |
| | **enrichment** | 1.05 | 1.08 | 1.04 | 1.01 | 1.06 | 0.96 | 1.01 | 1.07 |
| | ***p*-value** | 0.0684 | 0.0298 | 0.1025 | 0.3927 | 0.0761 | 0.8089 | 0.3293 | 0.0019 |
| **mRNA** | **count** | 510 | 370 | 529 | 777 | 363 | 296 | 789 | 1009 |
| | **enrichment** | 1.05 | 1.08 | 1.04 | 1.01 | 1.06 | 0.97 | 1.01 | 1.07 |
| | ***p*-value** | 0.0692 | 0.0323 | 0.1045 | 0.3704 | 0.0809 | 0.7996 | 0.3228 | 0.0023 |
| **exon** | **count** | 378 | 288 | 423 | 582 | 272 | 223 | 569 | 767 |
| | **enrichment** | 1.06 | 1.13 | 1.12 | 1.02 | 1.07 | 0.98 | 0.99 | 1.10 |
| | ***p*-value** | 0.0903 | 0.0066 | 0.0021 | 0.2958 | 0.0884 | 0.6337 | 0.6480 | 0.0014 |

| | | | | | | | | | |
|---|---|---|---|---|---|---|---|---|---|
| **CDS** | **count** | 360 | 279 | 410 | 563 | 256 | 214 | 546 | 739 |
| | **enrichment** | 1.06 | 1.16 | 1.15 | 1.05 | 1.06 | 0.99 | 1.00 | 1.12 |
| | ***p*-value** | 0.0885 | 0.0023 | 0.0008 | 0.1591 | 0.1288 | 0.5577 | 0.5153 | 0.0004 |
| **5' UTR** | **count** | 142 | 107 | 153 | 218 | 94 | 80 | 217 | 270 |
| | **enrichment** | 1.14 | 1.21 | 1.17 | 1.11 | 1.07 | 1.02 | 1.09 | 1.11 |
| | ***p*-value** | 0.0535 | 0.0244 | 0.0253 | 0.0802 | 0.2633 | 0.4512 | 0.0966 | 0.0348 |
| **3' UTR** | **count** | 117 | 88 | 143 | 178 | 89 | 72 | 161 | 242 |
| | **enrichment** | 1.10 | 1.17 | 1.28 | 1.05 | 1.18 | 1.07 | 0.94 | 1.17 |
| | ***p*-value** | 0.1448 | 0.0834 | 0.0015 | 0.2652 | 0.0557 | 0.2863 | 0.7917 | 0.0051 |

Significant values (*i.e.* $p < 0.05$) underlined.

**Table S5.** Enrichments of selective sweeps within groupings of insecticide resistance genes for each empirical mosquito population dataset.

| | | AOM | BFM | BFS | CMS | GAS | GNS | GWA | UGS |
|---|---|---|---|---|---|---|---|---|---|
| | | ***all four sweep states*** | | | | | | | |
| metabolism | count | 67 | 58 | 60 | 88 | 69 | 57 | 75 | 95 |
| | enrichment | 1.57 | 2.16 | 1.50 | 1.65 | 1.25 | 1.61 | 1.35 | 1.57 |
| | *p*-value | 0.0011 | ≤0.0001 | 0.0048 | ≤0.0001 | 0.0464 | 0.0018 | 0.0076 | 0.0001 |
| target site | count | 9 | 9 | 5 | 7 | 7 | 7 | 8 | 9 |
| | enrichment | 1.95 | 2.90 | 1.16 | 1.38 | 1.12 | 1.69 | 1.28 | 1.44 |
| | *p*-value | 0.0132 | 0.0007 | 0.4434 | 0.2087 | 0.4439 | 0.0823 | 0.2509 | 0.1096 |
| behavior | count | 18 | 19 | 15 | 20 | 20 | 18 | 18 | 20 |
| | enrichment | 1.73 | 2.63 | 1.42 | 1.51 | 1.37 | 2.04 | 1.24 | 1.26 |
| | *p*-value | 0.0507 | 0.0039 | 0.1619 | 0.1041 | 0.1518 | 0.0187 | 0.2480 | 0.2231 |
| cuticular | count | 2 | 1 | 2 | 6 | 5 | 2 | 5 | 7 |
| | enrichment | 0.82 | 0.60 | 0.87 | 1.77 | 1.49 | 1.02 | 1.62 | 1.94 |
| | *p*-value | 0.7046 | 0.8049 | 0.6813 | 0.1215 | 0.2408 | 0.5782 | 0.1921 | 0.0603 |
| | | ***completed hard*** | | | | | | | |
| metabolism | count | 0 | 0 | 2 | 0 | 0 | 0 | 0 | 1 |
| | enrichment | 0.00 | 0.00 | 26.63 | 0.00 | 0.00 | 0.00 | 0.00 | 8.64 |
| | *p*-value | 1.0000 | 1.0000 | 0.0121 | 1.0000 | 1.0000 | 1.0000 | 1.0000 | 0.0932 |
| target site | count | 0 | 0 | 0 | 0 | 0 | 0 | 0 | 0 |
| | enrichment | 0.00 | 0.00 | 0.00 | 0.00 | 0.00 | 0.00 | 0.00 | 0.00 |
| | *p*-value | 1.0000 | 1.0000 | 1.0000 | 1.0000 | 1.0000 | 1.0000 | 1.0000 | 1.0000 |
| behavior | count | 0 | 0 | 2 | 0 | 0 | 1 | 2 | 3 |
| | enrichment | 0.00 | 0.00 | 80.00 | 0.00 | 0.00 | 68.97 | 45.56 | 94.64 |
| | *p*-value | 1.0000 | 1.0000 | 0.0047 | 1.0000 | 1.0000 | 0.0109 | 0.0076 | 0.0020 |
| cuticular | count | 0 | 0 | 0 | 0 | 0 | 0 | 0 | 0 |
| | enrichment | 0.00 | 0.00 | 0.00 | 0.00 | 0.00 | 0.00 | 0.00 | 0.00 |
| | *p*-value | 1.0000 | 1.0000 | 1.0000 | 1.0000 | 1.0000 | 1.0000 | 1.0000 | 1.0000 |
| | | ***completed soft*** | | | | | | | |
| metabolism | count | 32 | 23 | 18 | 23 | 23 | 32 | 45 | 33 |
| | enrichment | 1.93 | 2.68 | 2.62 | 2.12 | 1.78 | 1.97 | 2.13 | 1.87 |
| | *p*-value | 0.0046 | 0.0008 | 0.0038 | 0.0036 | 0.0179 | 0.0021 | ≤0.0001 | 0.0053 |
| target site | count | 5 | 5 | 0 | 3 | 4 | 5 | 5 | 3 |
| | enrichment | 2.59 | 4.51 | 0.00 | 2.26 | 2.41 | 2.45 | 1.85 | 1.42 |
| | *p*-value | 0.0298 | 0.0025 | 1.0000 | 0.1322 | 0.0701 | 0.0390 | 0.1085 | 0.3543 |
| behavior | count | 11 | 4 | 2 | 10 | 12 | 9 | 7 | 9 |
| | enrichment | 2.79 | 1.79 | 1.13 | 3.91 | 3.89 | 2.29 | 1.29 | 1.98 |
| | *p*-value | 0.0180 | 0.2175 | 0.4666 | 0.0040 | 0.0025 | 0.0512 | 0.3263 | 0.0908 |
| cuticular | count | 1 | 0 | 0 | 0 | 1 | 2 | 5 | 4 |
| | enrichment | 1.06 | 0.00 | 0.00 | 0.00 | 1.54 | 2.24 | 4.42 | 3.84 |
| | *p*-value | 0.5901 | 1.0000 | 1.0000 | 1.0000 | 0.4597 | 0.2233 | 0.0075 | 0.0307 |
| | | ***partial hard*** | | | | | | | |

| | | | | | | | | | |
|---|---|---|---|---|---|---|---|---|---|
| **metabolism** | **count** | 11 | 8 | 18 | 36 | 32 | 10 | 8 | 24 |
| | **enrichment** | 1.40 | 1.77 | 1.39 | 2.58 | 1.02 | 1.27 | 1.16 | 3.27 |
| | ***p*-value** | 0.1944 | 0.1293 | 0.1453 | 0.0002 | 0.4736 | 0.2850 | 0.3799 | 0.0001 |
| **target site** | **count** | 0 | 0 | 1 | 4 | 3 | 1 | 0 | 1 |
| | **enrichment** | 0.00 | 0.00 | 0.66 | 2.70 | 0.80 | 1.11 | 0.00 | 1.07 |
| | ***p*-value** | 1.0000 | 1.0000 | 0.8033 | 0.0479 | 0.7734 | 0.6117 | 1.0000 | 0.6268 |
| **behavior** | **count** | 1 | 2 | 2 | 1 | 8 | 1 | 1 | 0 |
| | **enrichment** | 0.52 | 1.50 | 0.59 | 0.27 | 0.93 | 0.50 | 0.62 | 0.00 |
| | ***p*-value** | 0.7638 | 0.3395 | 0.7553 | 0.9470 | 0.5637 | 0.7603 | 0.6956 | 1.0000 |
| **cuticular** | **count** | 0 | 1 | 0 | 1 | 5 | 0 | 0 | 1 |
| | **enrichment** | 0.00 | 4.10 | 0.00 | 1.43 | 2.37 | 0.00 | 0.00 | 1.95 |
| | ***p*-value** | 1.0000 | 0.2027 | 1.0000 | 0.4852 | 0.0684 | 1.0000 | 1.0000 | 0.3747 |
| | | ***partial soft*** | | | | | | | |
| **metabolism** | **count** | 25 | 27 | 25 | 31 | 19 | 16 | 27 | 45 |
| | **enrichment** | 1.22 | 1.86 | 1.14 | 1.04 | 1.30 | 1.22 | 0.86 | 1.15 |
| | ***p*-value** | 0.2060 | 0.0095 | 0.2976 | 0.4337 | 0.1829 | 0.2658 | 0.7766 | 0.2125 |
| **target site** | **count** | 4 | 4 | 4 | 2 | 0 | 2 | 3 | 5 |
| | **enrichment** | 1.59 | 2.18 | 1.55 | 0.68 | 0.00 | 1.15 | 0.79 | 1.15 |
| | ***p*-value** | 0.2227 | 0.0966 | 0.2479 | 0.8377 | 1.0000 | 0.5334 | 0.7865 | 0.4586 |
| **behavior** | **count** | 6 | 13 | 9 | 9 | 2 | 7 | 9 | 10 |
| | **enrichment** | 1.19 | 3.36 | 1.56 | 1.27 | 0.55 | 2.23 | 1.11 | 0.95 |
| | ***p*-value** | 0.3837 | 0.0043 | 0.1779 | 0.3076 | 0.7908 | 0.0840 | 0.4277 | 0.5751 |
| **cuticular** | **count** | 1 | 0 | 2 | 5 | 0 | 0 | 0 | 3 |
| | **enrichment** | 0.82 | 0.00 | 1.58 | 2.36 | 0.00 | 0.00 | 0.00 | 1.32 |
| | ***p*-value** | 0.6906 | 1.0000 | 0.3598 | 0.0728 | 1.0000 | 1.0000 | 1.0000 | 0.3991 |

Significant values (*i.e.* $p < 0.05$) underlined.
